# Zinc-induced folding and solution structure of the eponymous novel zinc finger from the ZC4H2 protein

**DOI:** 10.1101/2025.06.19.660640

**Authors:** Rilee E. Harris, Antonio J. Rua, Andrei T. Alexandrescu

**Author notes:** CORRESPONDING AUTHOR: Andrei T. Alexandrescu. **ABBREVIATIONS:** AF3, Alpha Fold 3; CD, circular dichroism; DOSY, diffusion ordered spectroscopy; DSS, 3-(Trimethylsilyl)propane-1-sulfonic acid; EGTA, ethylene glycol-bis(β-aminoethyl ether)-*N*,*N*,*N*′,*N*′-tetraacetic acid; HSQC, heteronuclear single quantum correlation; LMCT, ligand-to-metal charge transfer; NLS, nuclear localization signal; NMR, nuclear magnetic resonance, NOE, nuclear Overhauser effect; NOESY, nuclear Overhauser effect spectroscopy; PDB, protein data bank; pLDDT, predicted local distance difference test; RMSD, root mean square deviation; SD, standard deviation; TCEP, tris(2-carboxyethyl)phosphine; TOCSY, total correlation spectroscopy; UV-Vis, ultraviolet-visible spectrophotometry; ZARD, ZC4H2-associated rare disorders; ZC4H2-ZL, long fragment comprised of residues 186-212 of the ZC4H2 protein; ZC4H2-ZS, short fragment comprised of residues 188-207 of the ZC4H2 protein; ZnF, zinc finger.

## Abstract

The *ZC4H2* gene is the site of congenital mutations linked to neurodevelopmental and musculoskeletal pathologies collectively termed ZARD (ZC4H2-Associated Rare Disorders). ZC4H2 consists of a coiled coil, and a single novel zinc finger with four cysteines and two histidines from which the protein gets its name. Alpha Fold 3 confidently predicts a structure for the zinc finger but also for similarly sized random sequences, providing equivocal information on its folding status. We show using a synthetic peptide fragment that the zinc finger of ZC4H2 is genuine, and folds around zinc ion with picomolar affinity. NMR pH titration of histidines and UV-Vis of a cobalt complex of the peptide indicate its four cysteines coordinate zinc while two histidines do not participate in binding. The experimental NMR structure of the zinc finger has a novel structural motif similar to RANBP2 zinc fingers, in which two orthogonal hairpins each contribute two cysteines to coordinate zinc. Most of the nine ZARD mutations that occur in the ZC4H2 zinc finger likely perturb this structure. While the ZC4H2 zinc finger shares the folding motif and cysteine-ligand spacing of the RANBP2 family, it is missing key substrate-binding residues. Unlike the NZF branch of the RANBP2 family, the ZC4H2 zinc finger does not bind ubiquitin. Since the ZC4H2 zinc finger occurs in a single copy it is also unlikely to bind DNA. Based on sequence homology to the VAB-23 protein, the ZC4H2 zinc finger may bind RNA of a currently undetermined sequence or have alternative unprecedented functions.

## 1. INTRODUCTION

ZC4H2 is a protein encoded by a gene located on the X-chromosome with largely unknown functions. Mutations in the *ZC4H2* gene, primarily expressed in the brain and central nervous system during embryonic development, are associated with central and peripheral nervous system neurodevelopmental disorders and musculoskeletal diseases collectively called ZARD [1–3]. The most common of these, Wieacker-Wolff syndrome (WRFF), is an X-linked recessive arthrogryposis multiplex congenita disorder, with manifestations including muscular atrophy, neurodevelopmental impairment, ocular defects, and respiratory problems [1, 4–7].

Some 90 interaction partners are listed for ZC4H2 in the IntAct database [8], with most of these identified from co-immunoprecipitation experiments. ZC4H2 is thought to bind to proteins in the SMAD pathway affecting bone morphogenic protein (BMP) signaling [9], which together with transforming growth factor-β regulates insulin transcription in pancreatic β-cells. These pathways have recently been implicated in a WRWF patient with hyperinsulemic hypoglycemia [10]. Another potential interaction target of ZC4H2 is the transient receptor potential cation channel subfamily V member 4 (TRPV4), a calcium ion channel that regulates cellular osmotic pressure. ZC4H2 is thought to bind to the cytosolic N-terminus of TRPV4, leading to increased basal activity and Ca^2+^ responses, together with increased channel turnover at the plasma membrane [11]. Notably, symptoms of ZARD resemble those of TRPV4-pathy, such as arthrogryposis, distal muscle weakness, club foot, and camptodactyly [12].

Finally, ZC2H2 has an intriguing possible role in proteostasis, through modulation of the Sonic Hedgehog (Shh) signaling pathway that governs embryonic spinal cord patterning and dendrite formation through glioma-associated oncogene (Gli) transcription factors [13]. ZC4H2 regulates the ubiquitination of RING finger protein 220 (RNF220), an E3 ubiquitin ligase enzyme that by attaching the small 76-a.a. ubiquitin protein, epigenetically controls Gli expression gradients and effectively serves as an Shh signaling enhancer [14–16]. Interactions between ZC4H2 and RNF220 have been implicated in the development of noradrenic neurons [15]. ZC4H2 also interacts with the E3 ubiquitin ligase RLIM, for which it itself is a substrate [17]. In partnership with the E3 ubiquitin ligases RNF220 and RLIM, ZC4H2 is poised to control protein levels post-translationally in several pathways involved in neural development. The interactions with the E3 ubiquitin ligases could form the basis of ZC4H2 dysfunction in as much as mutations in RNF220 and RLIM lead to similar neurological pathology phenotypes as ZARD mutations [14, 15]. To date there is no structural information on any protein interactions involving ZC4H2, or even on what parts of the protein could be involved in binding. Thus, structural studies are sorely needed for a mechanistic understanding of ZC4H2 [17].

The 224 a.a. human ZC4H2 protein (UniProt accession number Q9NQZ6) is predicted to consist of an α-helical coiled coil domain (residues 11-104) and a single putative novel ZnF domain (residues 189-206), from which the protein derives its name (Fig. 1A). Immediately following the ZnF, is a putative nuclear localization signal (NLS) thought to run between residues 207-224 [18]. The *ZC4H2* gene was first identified in a human brain gene sequencing project, and initially named KIAA1166 [19]. Subsequently, the protein was found to be a human liver cancer antigen, named HCA127 [20]. The designation of ZC4H2 as a ZnF protein is due to homology between HCA127 and an at the time novel sequence motif in the *C. elegans* epidermal morphogenesis regulator VAB-23 [21] described as a “putative C4H2 ZnF”. Fifteen years after the annotation of ZC4H2 [21], the ‘putative’ qualifier was dropped, but there is no published evidence that the sole globular domain in ZC4H2 or any of its homologs bind zinc or are functional ZnFs.

**Figure 1.**
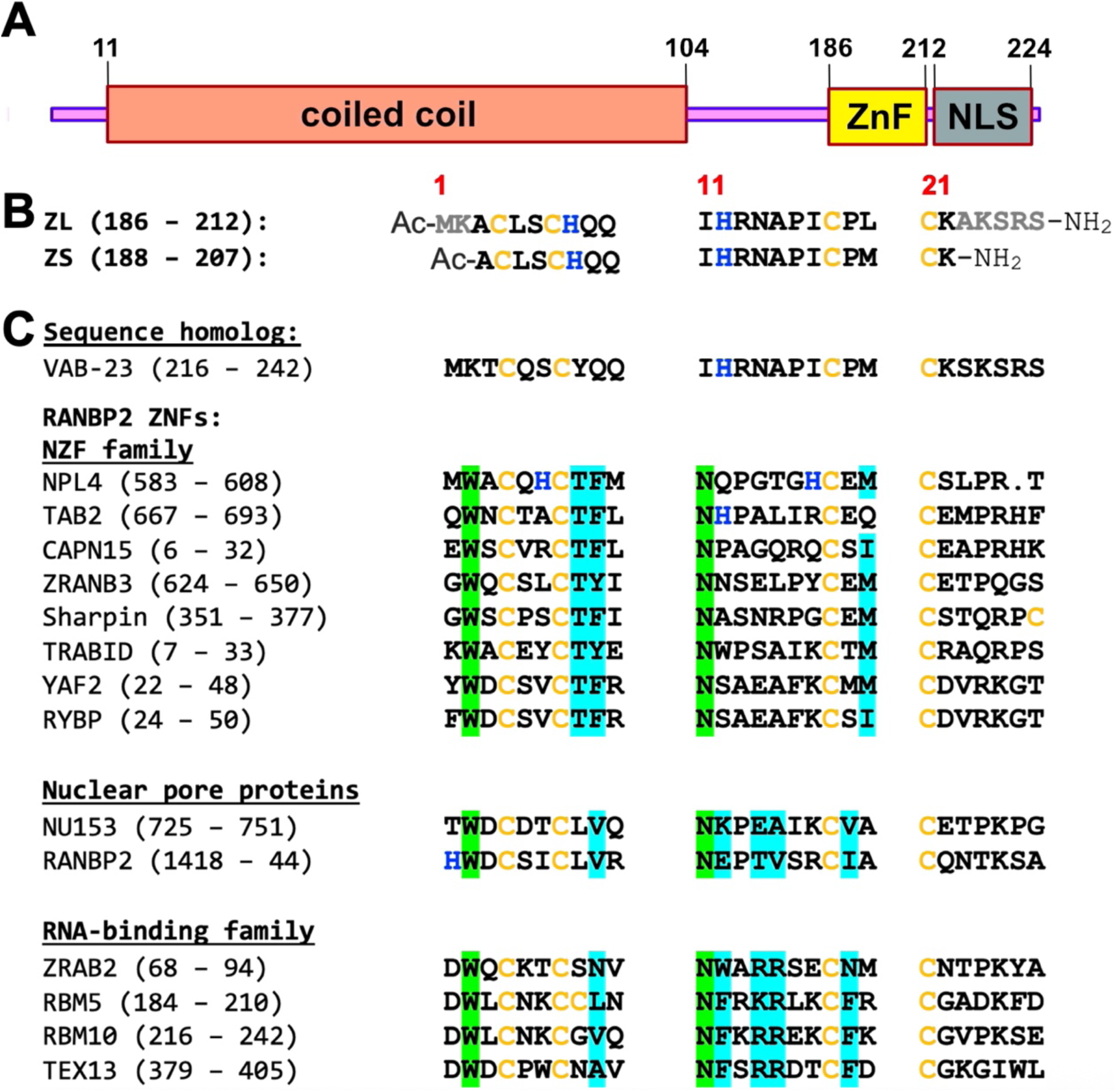
Domain organization and sequence of the ZnF from ZC4H2. (**A**) The human ZC4H2 protein consist of a coiled coil and a sole C-terminal ZnF domain that is the subject of this work. A putative nuclear localization signal (NLS) running from 207-224, overlaps with the C-terminus of the ZnF domain. (**B**) Sequence of the ZC4H2 ZnF. Initially we studied a short 20 aa fragment (188-207), ZS. Although ZS binds Zn^2+^, it gives NMR spectra similar to unfolded proteins. A longer 27 aa fragment (186-212), ZL, gives excellent NMR spectra and was used for structural studies. In this work, we use the 1-27 numbering scheme for the ZnF shown in red. (**C**) Sequences homologs of which *C. elegans* VAB-23 is the closest. The family of RANBP2 ZnFs have a C4-ligand spacing and structure similar to ZC4H2-ZL. The RANBP2 NZF subfamily typically function as ubiquitin-binding modules in ubiquitination pathways [55, 76]. NU153 and RANBP2 are nuclear pore proteins with protein-binding and possible DNA-binding functions [78, 86]. A third RANBP2 subfamily has RNA-binding functions [54]. All sequences except VAB-23 are for human proteins and only sequences with a two-residue spacing between the first two cysteines are shown. Cysteines and histidines are colored yellow and blue, respectively. Residues highlighted in green are conserved in all RANBP2 homologs but not ZC4H2 or VAB-23. Residues highlighted in cyan are involved in ubiquitin-binding in the NZF subfamily [76, 87] and nucleotide-binding in the RNA-binding subfamily [54].

Zinc fingers (ZnFs) are small 20-60 residue structural domains that fold upon binding one or more divalent zinc ions (Zn^2+^) [22–27]. About 3-5% of human proteins have ZnF domains, making this the most represented fold-family in the human genome [23, 28]. Because structures are largely stabilized by metal-binding, ZnFs are more structurally diverse than typical protein folds [27, 29, 30]. Over 50 ZnF subfamilies are known with cysteine (C) and histidine (H) Zn^2+^-ligand combinations such as CCHH, CCCH, CCHC, CCCC [26]. The CCHH DNA-binding family is the best characterized, occurring in about half of all transcription factors [23, 31]. While the CCHH ZnF family has a prototypical ββα fold (PDB: 1ZNF), other ZnF families have all-α (PDB: 1F81), all-β (PDB: 1K81), or folds lacking regular secondary structure altogether (PDB: 1PXE) [32]. The ZC4H2 domain does not have significant sequence homology to other sequence families, thus the Prosite database (accession ID PDOC51896) classifies it as a distinct type of ZnF with no known structural representatives.

NMR is ideally suited for determining the folding status and structure of ZnF domains. While it has been stated that cryoEM is the only structure determination technique worth investing in [33], typical ZnF domains at only 20-60 residues are too small for cryoEM. In multi-domain proteins, ZnFs are often separated from other domains by flexible linkers, so that their unconstrained rotational flexibility would lead to averaging of their electron densities into the noise. An exception is if the orientation of the ZnF domain were restricted by binding to other molecules, but it is unclear at this stage which, if any part of ZC4H2 binds other proteins. We found it particularly effective to use synthetic peptides to characterize ZnF domains, as this avoids the need for recombinant expression [34–37]. Here, we use synthetic peptide fragments to characterize the folding, metal binding, and structure of the novel ZnF domain from ZC4H2. We employ NMR and CD spectroscopy to show the ZC4H2 domain is a genuine ZnF that binds Zn^2+^ with high affinity. Because the ZnF has multiple Cys and His residues that are potential ligands for Zn^2+^, we use NMR pH titration of histidine residues and UV-vis spectroscopy of a Co^2+^ complex to identify the four residues that chelate the metal. Having established the coordination site, we determine the NMR structure of the ZnF and compare it to functional families that share the same metal-ligand spacing. Finally, we show evidence using two different sized peptide fragments, that Zn^2+^ chelation precedes the formation of stable tertiary structure in the domain.

## 2. MATERIALS AND METHODS

### 2.1 Samples

Fragments of human ZC4H2 (Fig. 1B) were made by solid phase peptide synthesis. The 27-residue peptide called ZL, corresponding to residues 186-212 was prepared to 97% HPLC purity by Biomatik (Kitchener, Canada). A shorter 20-residue peptide ZS, comprised of residues 188-207 was made to 90% purity by AAPPTec (Louisville, KY). Both peptides had acetylated N-termini and amidated C-termini to avoid the introduction of charges from free ends, thus better mimicking the fragments in the context of the intact protein. Mass spectrometry of both peptides gave values within 2 Da of the theoretical masses of 2,277 Da for ZS and 3,066 Da for ZL. Peptide concentrations were determined using the BCA assay [38]. ZnCl_2_ (anhydrous, purity ≥ 98%), CoCl_2_•6H_2_O (purity ≥ 99%), and all other reagents were from Sigma (St. Louis, MO).

### 2.2 NMR spectroscopy

To investigate zinc binding by NMR we looked at 1.6 mM samples of the ZS and ZL peptides in the absence or presence of equimolar ZnCl_2_, at pH 6.0 and a temperature of 10 C. The samples had no other added buffers or salts. For NMR assignments and structure determination we used two samples of the longer ZL peptide in the presence of equimolar ZnCl_2_. All of these experiments were done at 10 C, as temperatures above 25 C led to NMR line broadening and loss of amide proton signals. A 1.6 mM sample of ZL-Zn^2+^ in 90% H_2_O/10% D_2_O at pH 6.0 was used to collect 2D TOCSY (70 ms mixing time), NOESY (150 ms mixing time) and ^1^H-^15^N HSQC (natural abundance) experiments in protonated solvent on a Bruker NEO 800 MHz spectrometer. A second 0.9 mM sample of ZL-Zn^2+^ in 99.8% D2O at pD* 5.9 was used to collect 2D TOCSY (70 ms mixing time), NOESY (50 ms mixing time) and multiplicity-edited ^1^H-^13^C HSQC (natural abundance) experiments in deuterated solvent on a Bruker NEO 600 MHz spectrometer. Both spectrometers were equipped with cryogenic probes. Internal DSS (2,2-dimethyl-2-silapentane-5-sulfonate) was used for ^1^H and ^13^C referencing, while an indirect 3 (^15^N/^1^H) ratio of 0.101329118 was used for ^15^N referencing [39]. We obtained > 80% sidechain assignments and > 92% backbone assignments for ^1^H, ^13^Cα and ^15^N resonances but the ^15^N assignments should be considered tentative particularly for weak resonances, due to the low sensitivity of the ^1^H-^15^N HSQC spectrum at natural isotope abundance.

Additional NMR experiments were done to study the biophysical properties of ZL-Zn^2+^. A pH titration of 0.9 mM ZL-Zn^2+^ in D_2_O at a temperature of 10 C was used to investigate if histidines were involved in Zn^2+^-binding. The pH data were fit to a modified Henderson-Hasselbach equation to obtain histidine p*K*_a_ values [40]. DOSY spectroscopy [41] to verify the monomeric state of ZL-Zn^2+^ was done on a 0.3 mM sample in D_2_O, at pD* 6.9 and a temperature of 25 C. A temperature titration to assess the thermal stability of ZL-Zn^2+^ was done on a 0.3 mM sample in D_2_O, at a pD* of 6.9. Data up to 50 C were collected on a 600 MHz Varian Inova spectrometer but data at higher temperatures were collected on a 500 MHz Bruker Avance instrument, since our Varian probe did not allow work above 50 C. NMR data were processed with iNMR (http://www.inmr.net/) and analyses were performed with the program CcpNmr Analysis 2.5.2 [42].

### 2.3 CD spectroscopy

CD experiments were performed on an Applied Photophysics Chirascan V100 Spectrometer (Surrey, UK) using a 1 mm cuvette path-length, a 1 nm bandwidth, a 1 nm scan step size, and 5 s/point data averaging. ZS and ZL sample concentrations were 40 and 80 µM, respectively. The samples were in 5 mM HEPES buffer (pH 6.8 to 6.9), containing 0.2 mM of the reducing agent TCEP (tris(2-carboxyethyl)phosphine) to prevent cysteine disulfide formation. To measure metal binding affinity, ZnCl_2_ titrations at a temperature of ∼22 C were done in the presence of 10 mM competitive chelator EGTA [36, 43]. The *K*_d_ was calculated from the EGTΑ competition experiments as previously described [36].

### 2.4 UV-visible spectroscopy of Co^2+^ complexes

Spectra as a function of CoCl_2_ were recorded from 200-800 nm on an Ultrospec 8000 double-beam spectrophotometer (Thermo Fisher) using 500 µL samples in 750 µL cuvettes with a 1 cm pathlength at a temperature of 25 C. The ZS and ZL sample concentrations were 390 and 200 µM, respectively. Samples were in 10 mM TRIS buffered to pH 7.0, with 0.5 mM TCEP. Samples were blanked against cuvettes containing buffer alone.

### 2.5 Alpha Fold 3 simulations of random sequences

Five random sequences each, for protein lengths between 20 and 150 in increments of 10 residues, were generated with the random sequence server of EXPASY (https://web.expasy.org/randseq/). The sequences were submitted to the Alpha Fold 3 (AF3) server (https://alphafoldserver.com/) to generate structure predictions [44]. For random sequences matching the cysteine and histidine composition of the ZnF from ZC4H2, we generated 25 random sequences with the sequence length set to 27 residues, and specified cysteine and histidine compositions of 14.8% and 7.4%, respectively. The latter sequences were submitted to AF3 for prediction with inclusion of a single Zn^2+^ ion. All of the random sequences used for this work are available from the corresponding author upon request.

### 2.6 NMR structure calculations

Dihedral angle restraints were calculated from assigned chemical shifts using the program TALOS-N [45]. Distance restraints from NOESY spectra were set to three upper ranges of 3.0, 4.0 and 5.0 Å respectively, based on peak intensities. The lower bound was uniformly set to 1.8 Å. Pseudo-atom upper bound corrections of 1.0, 2.0, 2.4 and 1.5 Å were included for protons with ambiguous methylene, aromatic ring, prochiral methyl, and methyl group NMR signals, respectively [46]. Five hydrogen bond restraints (four β-hairpin and one α-helix) were included based on the secondary structure consensus, and proximity of hydrogen bond donors and acceptor atoms in initial NMR structures excluding hydrogen bonds. The Zn^2+^ atom and ligands were restrained using distance bounds of 2.33-2.37 Å for Zn^2+^-Sψ and = 3.25-3.51 Å for Zn^2+^-Cβ [35] for each of the four cysteines. An additional six restraints of 3.02-4.52 Å were included between each pair of cysteine Sψ atoms to enforce the tetrahedral geometry of the binding site [47]. The PdbStat program was used to remove redundant and structurally non-informative restraints [48].

NMR structure calculations were initially performed on the NMRbox platform [49] with the program X-Plor NIH v. 3.8, using the protein-4.0 parameter set [50]. Structures were calculated using distance geometry, followed by simulated annealing refinement using the X-plor script *prot_sa_refine_nogyr.inp* from the NESG site (https://nesgwiki.chem.buffalo.edu).

Once we obtained restraint sets without violations, we further water-refined the structures using the program ARIA 2.3.2 [51] on NMRbox. A set of 200 structures in explicit H_2_O solvent were calculated with the default iteration protocols of the ARIA program, from which the 20 structures with the lowest energies and no violations were chosen for deposition and analyses (Table S1).

### 2.7 Database accession codes

Chemical shifts for ZL-Zn^2+^ were deposited in the BioMagResBank under accession number 53214. Restraints and coordinates for NMR structures were deposited in the Protein Data Bank under accession ID 9P3Z.

## 3. RESULTS

### 3.1 ZC4H2 has a genuine ZNF with domain boundaries larger than those specified by UniProt

The UniProt database predicts the ZC4H2 protein is comprised of a coiled coil, followed by single C-terminal ZnF domain with a unique C-X2-C-H-X3-H-X5-C-X2-C sequence pattern (Fig. 1A,B). Since no members of the ZC4H2 homology family have yet been shown to be genuine ZnFs, we first wanted to see if the domain from the human protein folded in the presence of Zn^2+^. We initially studied a synthetic peptide fragment extended by one amino acid on either side of the 189-206 domain boundary specified by the UniProt database (accession code Q9NQZ6). We call this short 20 a.a. segment ZS. Although there are some differences between the 1D NMR spectra of the peptide with and without Zn^2+^ (Fig. 2A, ZS), these are small, and the amide proton NMR signals largely fall in the random coil region of the spectrum. The 2D TOCSY spectrum of ZS similarly has random coil-like chemical shifts, as well as NMR line broadening characteristic of dynamics on the µs-ms timescale (Fig. 2B, ZS-Zn^2+^). Despite the poor NMR spectra, the ZS fragment undergoes changes in CD spectrum characteristic of a folding transition with increasing Zn^2+^ concentration (Fig. 3A). Using a previously described assay measuring CD signal changes upon Zn^2+^ addition in the presence of the competitive chelator EGTA [37, 43], we calculate the ZS fragment binds Zn^2+^ with a *K*_d_ of 18 pM (Fig. 3C).

**Figure 2.**
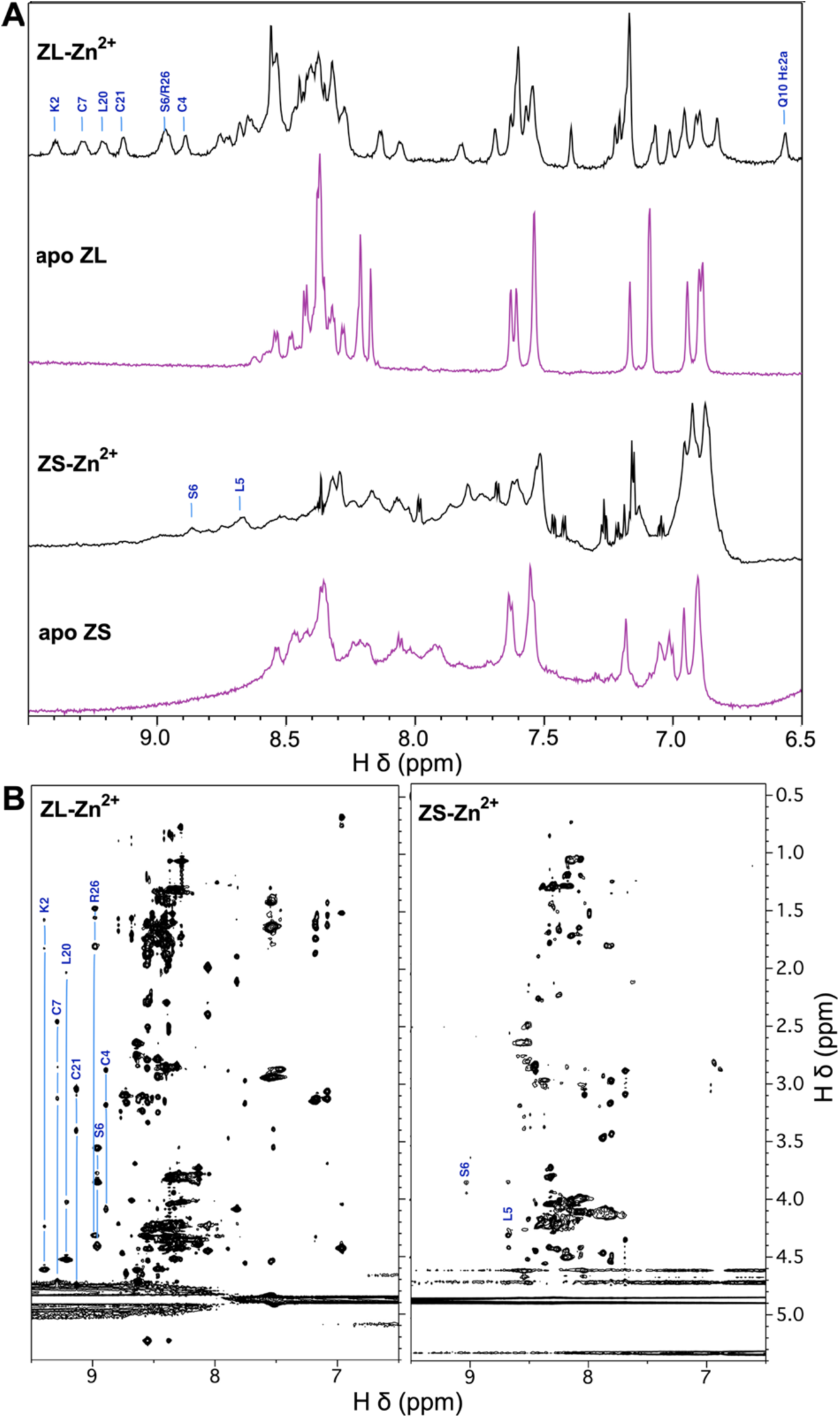
Comparisons of NMR spectra for the ZS and ZL peptide fragments. (**A**) 1D ^1^H-NMR spectra of ZS and ZL in the absence and presence of equimolar ZnCl_2_. (**B**) 2D-TOCSY (70 ms mixing time) of ZS and ZL both with equimolar Zn^2+^. NMR spectra were collected at 800 MHz on 1.6 mM peptide samples at pH 6.0 and a temperature of 10 C. Selected sequence-specific assignments are given for well-dispersed NMR signals based on the complete assignments for zinc-bound ZL (Fig. S2). Note the poorer NMR dispersion for ZS, which has random-coil-like chemical shift ranges (HN ∼8.7-7.7 ppm and Hα ∼4.6-4.0 ppm).

**Figure 3.**
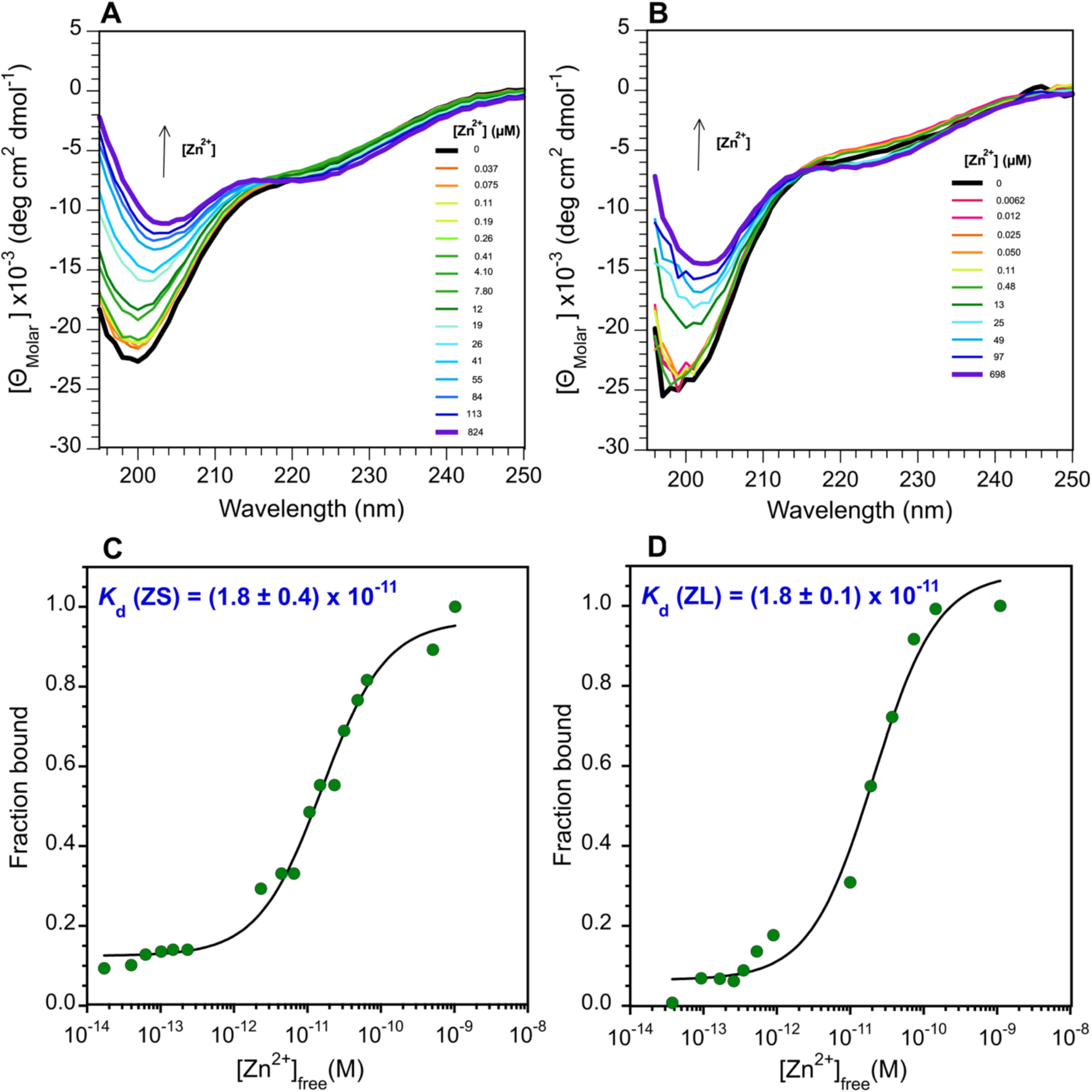
Zn^2+^ binding followed by CD spectroscopy. ZnCl_2_ titrations with peptide fragments ZS (**A**) and ZL (**B**). CD experiments were done at a temperature of 20 C on 80 µM peptide samples in 5 mM HEPES buffer pH 6.8-7.0, containing 0.2 mM of the reducing agent TCEP, 10 mM of the competitive chelator EGTA, and the indicated ZnCl_2_ concentrations. Fits of the binding data as previously described [36, 43], were used to obtain *K*_d_ values for Zn^2+^-binding to the ZS (**C**) and ZL (**D**) fragments. Reported *K*_d_ values are the mean ± SD from duplicate experiments on separate samples of each of the ZS and ZL fragments.

We suspected that the ZS fragment gave poor NMR spectra because its domain boundaries are too short, disturbing the structure of the ZnF. We consequently looked at a longer 27 a.a. fragment encompassing residues 188-207 of ZC4H2, called ZL. Compared to ZS, the ZL fragment extends the sequence by two residues at the N-terminus (Fig. 1B) that are involved the β-hairpin structure of ZnF homologs (Fig. 1C). We also extended the C-terminus with the stretch of polar residues AKSRS (Fig. 1B), expecting this would improve solubility. The ZL fragment shows several backbone and side chain amide protons outside the random coil region of the spectrum (labeled in Figs. 2A,B), and while the fragment lacks any aromatic residues other than two histidines, it gives excellent NMR chemical shift dispersion. DOSY spectra [41] show the Zn^2+^-bound ZL fragment is a monomer (Fig. S1).

Although NMR line broadening affects signals from ZL at temperatures above ∼25 C, NMR spectra improve at a temperature of 10 C. The good chemical shift dispersion of the ZL fragment (Fig. 2A, ZL-Zn^2+^) allowed us to obtain NMR assignments for the entire sequence (Fig. S2). Despite the superior NMR spectra for ZL, the changes in the CD spectrum of ZS with Zn^2+^ are similar to those for the shorter ZS fragment (Fig. 3A,B). Moreover, the *K*_d_ of 18 pM for Zn^2+^ for the ZL fragment is the same as that for the ZS fragment (Fig. 3C,D). This suggests the Zn^2+^ binding site as detected by CD is similar in the ZS and ZL fragments, but only the longer ZL fragment achieves a stable tertiary structure as monitored by NMR.

Compared to other ZnFs we recently studied that have thermal melting points > 80 C [34, 36, 37], ZC2H4-ZL undergoes a thermal unfolding transition by NMR at a much lower temperature of 32 ± 6 C (Fig. S3). Broadening of NMR resonances and loss of amide peaks at high temperatures is why we did all structural studies with ZC4H2-ZL at a temperature of 10 C. Although the changes in the NMR spectrum with temperature are completely reversible, the NMR signals from the thermally unfolded state are broad (Fig. S3, Uχ1 and Uδ2), suggesting NMR relaxation contributions from motion on µs-ms timescale typical of molten globules [52, 53], or indeed the ZS fragment (Fig. 2A). This is not the case with the protein unfolded by acidic pH, where the NMR signals from the unfolded state are sharper than those from the folded state (Fig. 4B, Uχ1 and Uδ2). These observations raise the possibility that the thermally unfolded state still has zinc bound. Of note, CD spectra of the samples at high temperatures are more similar to the zinc-bound than free peptide. We note the thermal unfolding transition observed by NMR occurs near physiological temperatures, but do not know if this has functional significance.

**Figure 4.**
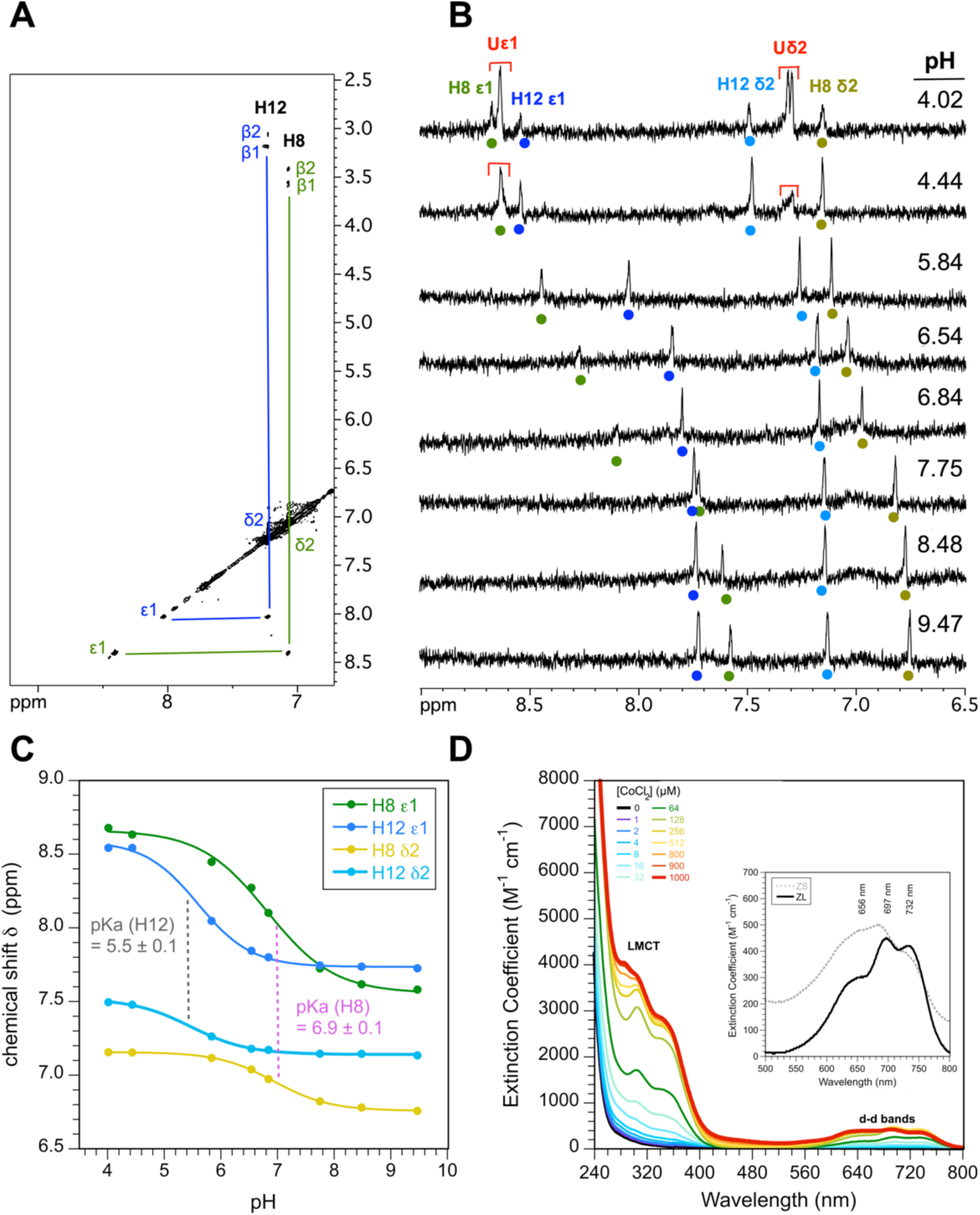
Identification of the four cysteines as metal ligands. (**A**) Assignments of histidine ring protons from ^4^J_H82-Hβ_ couplings in a 70 ms mixing time TOCSY recorded for ZL-Zn^2+^ in D_2_O. (**B**) pH titration of the H8 and H12 aromatic ring resonances. Note that the two histidines are the only aromatic residues in ZL. The fact that both histidines shift with pH indicates that they do not participate in metal coordination. The signals Uχ1 and U82 (red) are from the histidines in the acid-unfolded state. (**C**) Titration curves for the two histidines in the folded state and p*K*_a_ values (mean ± SD) calculated from the H82 and Hχ1 curves for each histidine. (**D**) Titration of ZL with CoCl_2_. The LMCT (315 nm) and d-d bands (500-800 nm) are consistent with coordination of the metal by four cysteines [59]. The inset compares the d-d band region for ZL (black line) and ZS (dotted gray line), both in the presence of 128 µM CoCl_2_. The similarity of the spectra indicates similar metal coordination sites for ZS and ZL.

### 3.2 The ZnF of ZC4H2 has a CCCC metal coordination sphere

NMR data for the ZL fragment appeared amenable to structure determination. However, ZnFs such as ZC4H2 with multiple potential metal ligands can confound the identification of metal-coordinating residues [36, 37]. The four cystines and two histidines in ZL can be arranged in 11 different coordination combinations to bind a single zinc atom (1 CCCC, 4 CCCH, and 6 CCHH). Considering a four cysteine coordination set, the C-X2-C-X10-C-X2-C sequence spacing in ZC4H2 is similar to the RANBP2 family [54, 55] with a C-X(2-4)-C-X10-C-X2-C sequence pattern (Fig. 1C) but also to the N-terminal part of the C6 ZnF from lysine-specific demethylase hairless (UniProt O43593, a.a. 600-625) which has a <u>C-X2-C-X10-C-X2-C</u>-X4-C-X2-C sequence pattern. The underlined portion of the sequence is the part similar to ZC4H2-ZnF. Other similar cysteine spacings are found in parts of ZnFs that bind two zinc ions, such as the first half of the MYND domain: <u>C-X2-C-X(7-11)-C-X2-C</u>-X5-C-X3-C-(X7,8)-H-(X3)-C (Uniprot Q8IYR2, a.a. 296-335) and the second half of ring fingers such as MGRN1: C-X2-C-X(9-39)-C-X(1-3)-H-X(2-3)<u>-C-X2-C-x10-C-X2-C</u> (Uniprot O60291, a.a. 278-317). We therefore felt an important prerequisite for structure determination was to first accurately identify the residues that participate in zinc coordination.

To this end we obtained NMR pH titration data for the ZL fragment. The rationale is that histidines bound to zinc should be invariant to changes in pH when they are bound to zinc [36, 56]. By contrast, histidines not involved in metal coordination should titrate freely with pH as they undergo their protonation-deprotonation equilibria. Starting from our complete backbone NMR assignments, we first assigned the aromatic ring protons of the two histidines through weak four-bond ^4^J_H82-Hβ#_ couplings observed in the 70 ms TOCSY spectrum for the sample in D_2_O (Fig. 4A). The data for the aromatic region of the ZL NMR spectrum (Fig. 4B) indicates both histidines titrate freely between pH 4.0 and 9.5, so that they are not involved in zinc coordination. Rather, the four cystines in ZC4H2 must form the zinc coordination site. Of the two histidines, H8 has a p*K*_a_ of 6.9 in the random coil range, whereas the p*K*_a_ of 5.5 for H12 is lowered by about 1 to 1.5 pH units compared to random coil values (Fig. 4C). The lowered p*K*_a_ of H12 is probably due to a repulsive electrostatic interaction with the positively charged guanidino group of R13. Repulsion between the positive charges of R13 and H12 would resist protonation, needing a larger proton concentration (lower pH) to ionize H12 [57, 58].

Metal coordination was further studied using a CoCl_2_ titration of the ZL fragment monitored by UV-Vis spectrophotometry (Fig. 4D). The Co^2+^ titration supports the coordination of a single metal ion by four cysteines. The ligand-metal-charge-transfer (LMCT) band at metal saturation has an χ_315_ = 4000 M^-1^ cm^-1^, consistent with four cysteines binding to the metal since each thiolate group is expected to contribute ∼ 1000 M^-1^ cm^-1^ [59]. The d-d transition bands between 500-800 nm are red-shifted, which is also consistent with thiol groups exclusively contributing to metal coordination [60, 61]. The inset compares the d-d band spectrum for the ZL and ZS fragments (Fig. 4D). Despite the differences in the quality of the NMR spectra of the two fragments, the d-d bands in the UV-Vis spectrum show they bind metal using a very similar tetrahedral CCCC-coordination mode.

### 3.3 Alpha Fold 3 predicts the ZnF structure with confidence but also similarly sized random sequences

The AF3 program [44] predicts a structure for the ZL fragment consisting of two orthogonal hairpins, with each of the hairpins contributing two of the cysteines to the zinc binding site (Fig. 5A). The structure is typical of the RANBP2 family of ZnFs [54, 55] and also occurs in iron-binding proteins such as rubredoxin. The hairpin motif, of which there are two in the model, has been called the ‘rubredoxin knuckle’ [55]. In the AF3 model, the four cysteines in the sequence coordinate Zn and the histidines do not participate, consistent with our results from pH titrations and UV-Vis spectroscopy of cobalt complexes (Fig. 4). Given that AF3 confidently predicts a structural model for the ZnF of ZC4H2, a reasonable question is whether it is worthwhile to calculate an experimental NMR structure, a process that is considerably more time-consuming and laborious than an AI-generated structure prediction. As outlined below we feel that even if the AF3 prediction is trustable, it is important to verify this experimentally for targets such as the ZnF of ZC4H2 that have no representative structures.

**Figure 5.**
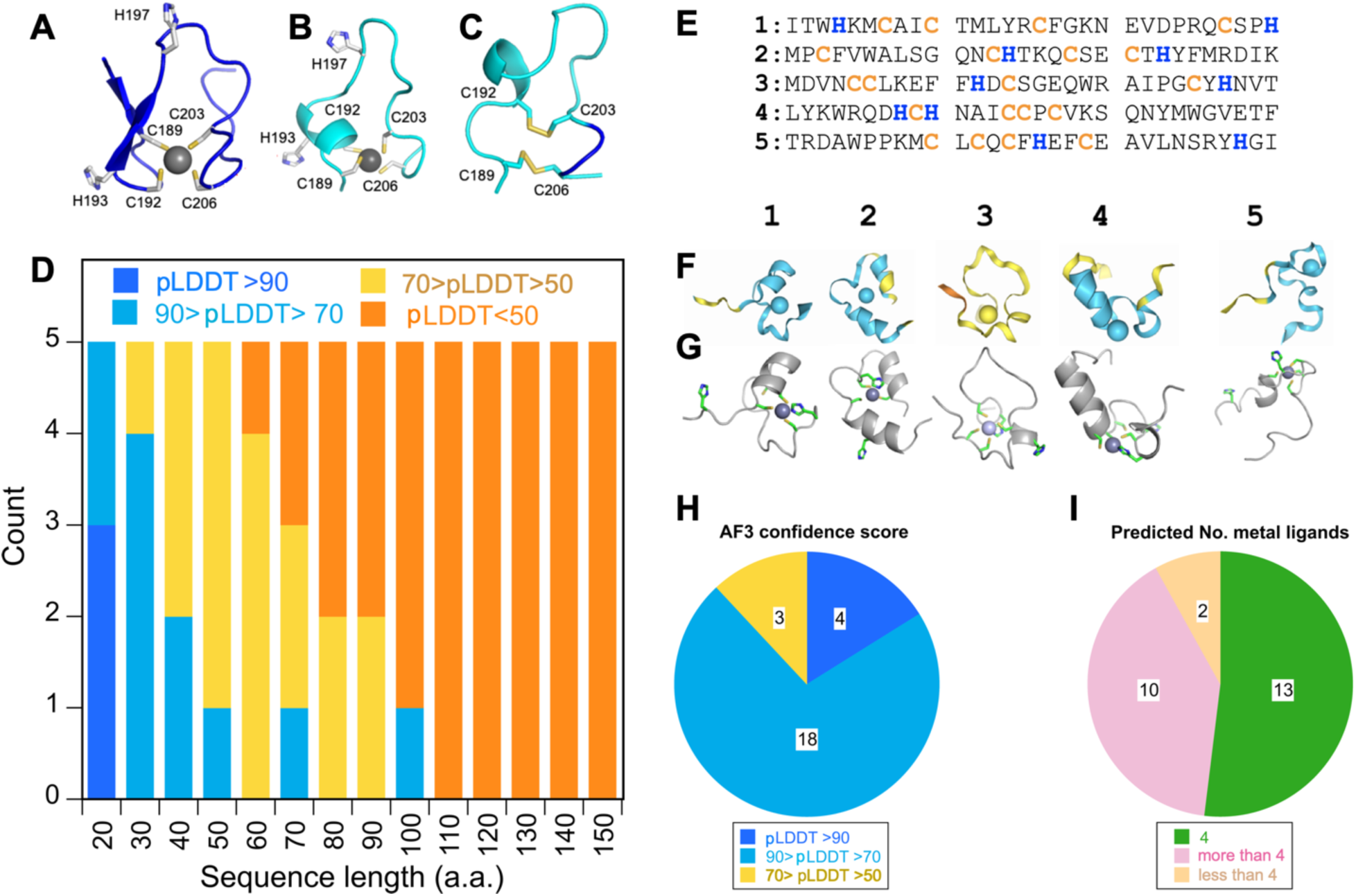
Alpha Fold 3 (AF3) predictions. **(A)** ZL fragment with Zn^2+^. **(B)** ZS fragment with Zn^2+^. Note the difference in metal coordination of C189 and C192 compared to ZL in (A). (**C**) The ZS fragment without metal is predicted to form disulfide bonds. (**D**) To test how AF3 confidence (LDDT score) depends on sequence length, we generated five random sequences for each of the specified lengths between 20 and 150 a.a. For random sequences shorter than 100 a.a., and particularly for those shorter than 50 a.a., AF3 gives high pLDTT scores (dark and light blue) that eventually decrease (yellow and orange) for the longer sequences between 100-150 a.a. (**E**) Five representative random sequences constrained to have a total of 30 a.a. with four cysteines (yellow) and two histidines (blue), the same distribution of metal-ligating residues as ZC4H2. (**F**) AF3 predictions of the five random sequences in (E), colored according to AF3 pLDDT confidence scores. Most of the random sequence predictions have the same confidence levels as the AF3 predictions for the ZNF from ZC4H2 in A-C. (**G**) AF3 predictions from panel F showing the cysteine and histidine residues. In all cases four Cys/His residues form a coordination site for Zn^2+^. To obtain better statistics we simulated a larger set of 25 random sequences, each with 27 residues like ZC4H2-ZL and 4 Cys and 2 His. The pLDDT confidence levels for the random sequence predictions (**H**) and the number of cystine and histidines bound to Zn^2+^ for the predicted structures (**I**).

While the two-hairpin motif is predicted for the ZL fragment (Fig. 5A), for the shorter ZS fragment AF3 predicts a loop structure in which both the individual hairpins are lost (Fig. 5B). Note that there is also a switch between the positions of the two zinc ligands C189 and C192 (relative to C203 and C206) in the two models (Fig. 5A,B). AF3 calculates a model similar model to that in Fig. 5A when the ZL prediction is done without zinc, but with the shorter ZS fragment a loop linked by two disulfide bonds is predicted in the absence of zinc (Fig. 5C). All of the models have light or dark blue colors corresponding to confident or very confident predictions (pLDDT scores better than 70). Yet there are differences in the predictions depending on the sizes of the protein fragments considered for the structural domains.

To gain further insight into the confidence levels of AF3 predictions for domains similarly sized to the ZnFs of ZC4H2, we tested how prediction confidence varied with increasing sequence length. We used the RandSeq server of ExPASy [62] to generate five random sequences for each polypeptide length between 20 and 150 residues, in increments of 10 amino acids (Fig. 5D). The majority of the AF3 predictions had substantial and realistic-looking secondary structures. For sequences longer than 100 residues, the pLDDT scores were predominantly below 50, corresponding to low-confidence predictions as expected for random amino acid sequences. For sequences shorter than 100 amino acid, however, the predictions increased in confidence with decreasing sequence length. The data in Fig. 5D consider the overall pLDDT score for the prediction. In nearly all groups of five predictions shorter than 100 residues, at least one of the five predictions had a segment with a pLDDT score better than 70, even when the protein had an overall lower score. These confident predictions could correspond to structured domains or subdomains in proteins that are otherwise unfolded, and this is not accounted for in the data of Fig. 5D. We are not entirely sure why AF3 confidently predicts apparent false positive structures for proteins shorter than ∼50 amino acids with random sequences. This could be due to the spurious ‘hallucination’ of structures for disordered regions, a noted limitation for the diffusion-based AI algorithm for AF3 [44]. Alternatively, the occurrence of high-confidence predictions for short random sequences could be a type of “Russian doll effect” [63], where shorter sequences are statistically more likely to match a sequence in the X-ray structure database used for AI training. Since AF3 cannot distinguish structured from disordered polypeptides, the occurrence of a reasonable match in the X-ray database raises the confidence of the prediction, even when the matching short segment would likely be unfolded outside the context of the larger protein.

We saw similar behavior with a set of 25 random sequences selected to match the 27-residue length and 4 cysteine plus 2 histidine content of the ZnF in ZC4H2. Five representatives of these random sequences are given in Fig. 5E and their corresponding AF3 predictions in Figs. 5F,G. Statistics pertaining to pLDDT confidence levels for the predictions, and to the number of Cys/His coordinating the zinc ion are summarized in Fig. 5H and Fig. 5I, respectively. Most of the 25 random sequences in the trial dataset are confidently predicted (88% pLDDT > 70) with four giving very confident predictions (16%, pLDDT > 90). These observations are consistent with the behavior of 30-residue polypeptides in the larger dataset of random sequences (Fig. 5D). Note that the observed pLDDT confidence scores for the random sequences are of the same order as those for the prediction of the actual sequence for the ZnF in ZC4H2 (Fig. 5A-C). Remarkably, the majority of ZC4H2-like random sequences (52%) have a combination of four Cys/His residues (green in Fig. 5G) within bonding distance of the zinc ion, with a further 40% having more than four Cys/His near the metal, and 8% having only three Cys/His in the coordination site. While rare, folded ZnFs with only three of four metal ligands coordinating zinc occur in nature [36]. Thus, the random ZnF-like sequences have a high chance to give realistic-looking zinc coordination sites in AF3 predictions (Fig. 5I).

Small proteins, including ZnF domains, are often viewed as trivial “low-hanging fruit” in current structural biology. However, it is becoming increasingly appreciated that there is a vast repertoire of thousands of biologically important mini and microproteins shorter than 50-100 amino acids [64, 65] who’s structural properties are largely unexplored. Given the ambiguities of AF3 predictions for such small proteins and their structural variability, exemplified by ZnFs that includes some 50 different types of structural motifs, these mini and microproteins could represent a future frontier for experimental structural biology. As funding priorities increasingly shift from curiosity-driven science to AI, technology, and for-profit research, these structures may unfortunately remain uncharted at least in the short term.

### 3.4 The NMR structure of the ZC4H2 is a variant of the RANBP2 ZnF fold

The experimental NMR structure of the ZL fragment corresponding to the ZF in ZC4H2 is shown in Figs. 6A,B. The structures are shown on a color ramp from the N-(blue) to the C-terminus (red). The folding motif has some similarity to the RANBP2 ZnF fold, where two orthogonal glutaredoxin knuckle β-hairpins each provide two cysteines to form a coordination cage for zinc [54, 55]. Conceptually, this fold replaces the α-helix in the classical ββα motif of CCH(H/C)-type ZnFs [23, 37] with a second β-hairpin to give a ββ-ββ structure. In the ZnF of ZC4H2 the second hairpin lacks canonical β-sheet secondary structure since the segment corresponding to the third β-strand has two proline residues that preclude β-sheet hydrogen bonding. Other ZnFs with glutaredoxin knuckles such as TAB2-NZF [66] and the RANBP2 ZnF from FUS [67] also do not have β-sheet structure for the second hairpin. A unique feature of the ZnF in ZC4H2, is the presence of a short one turn α-helix at the end of the second hairpin (Fig 6A). This proved difficult to identify as it is not predicted by AF3 (Fig. 5A) but is supported by α-helical chemical shift indices as well as short range NOEs characteristic of α-helical structure for residues C21-S25 at the C-terminus of ZL (Fig. S3). Consequently, the RMSD between the NMR structure and the AF3 model which probably derives its prediction from the large database of RANBP2 ZnFs in the PDB, is rather large at about 4 Å. Prediction are typically considered accurate when the RMSD is below a threshold of about 2.5 Å [68].

**Figure 6.**
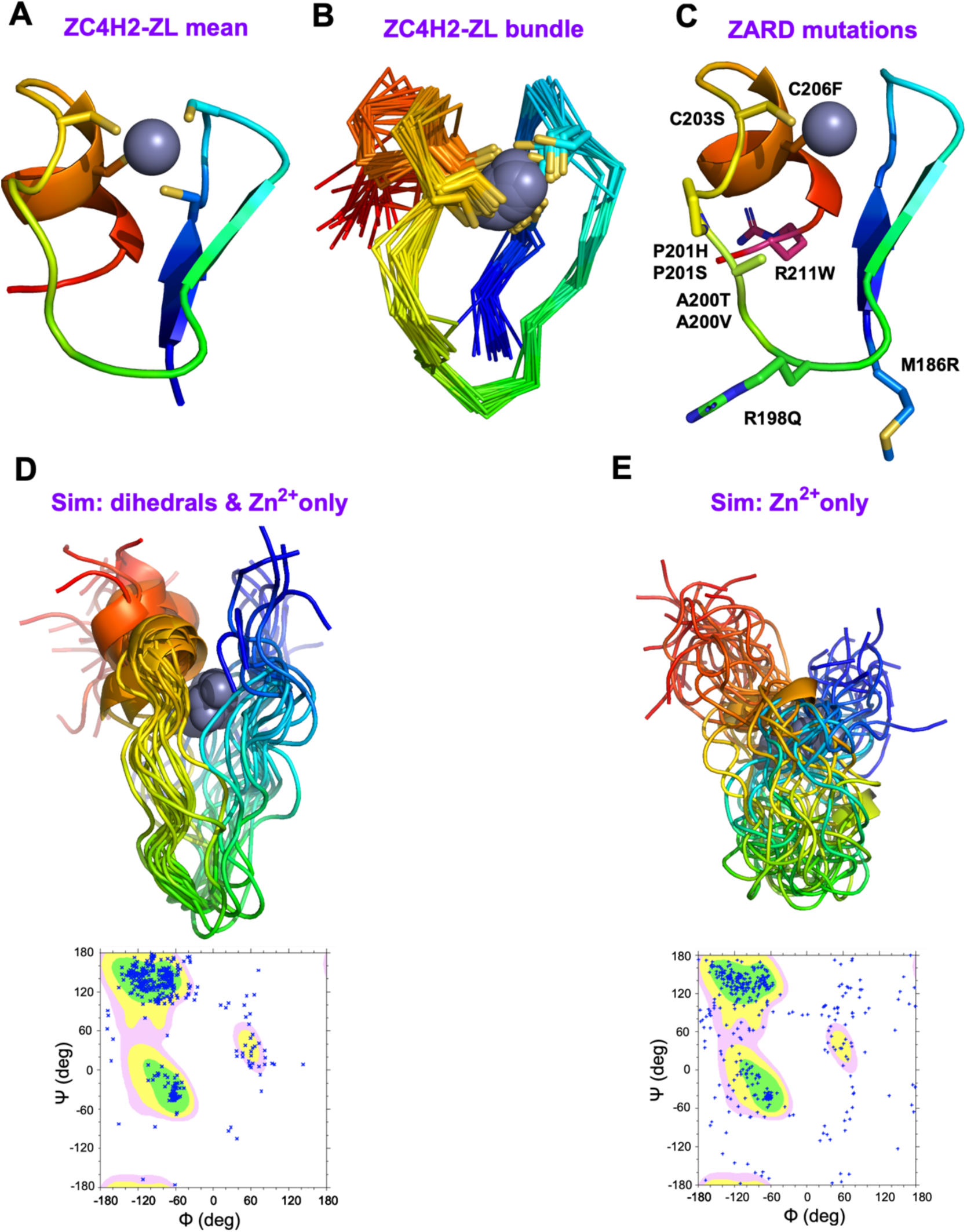
Structures of ZC4H2-ZL. The protein mainchain is shown on a color gradient from the N-term (blue) to the C-term (red). Sidechains of the four chelating cysteine residues are shown with sticks, and the zinc atoms as gray spheres. (**A**) NMR structure closest to the ensemble mean. (**B**) The NMR bundle. (**D**) Locations of nine known ZARD mutations in the ZnF, labeled according to the numbering scheme for the full-length ZC4H2 protein. (**D**) Simulated structures calculated with only dihedral and zinc ion restraints, and the associated Ramachandran plot. (**E**) Simulated structures calculated with only restraints to Zn^2+^ and the associated Ramachandran plot.

Although the short α-helix is present in most NMR conformers for the ZL fragment, its relative position is less well defined than the remainder of the structure (Fig. 5B). The chemical shift indices predict a gradient of increasing disorder for the last four residues at the C-terminus, with the last two residues giving chemical shift-predicted S^2^ order parameters below 0.5. Recall that we included the last four residues 24-KSRS-27 to improve the solubility of the ZL fragment. The highly polar character of these amino acids probably accounts for the disorder observed at the C-terminus of the NMR structure.

## DISCUSSION

### 4.1 Structural mapping of the eight ZARD mutations within the ZnF

Our work establishes that ZC4H2 has a genuine folded ZnF for which we determined the NMR structure. We can use our structure (Fig. 6C) to analyze the location of some nine known ZARD mutations distributed amongst seven sites in the ZnF domain: M186R, R198Q, A200T, A200V, P201H, P201S, C203S, C206F and R211W [2, 3, 10]. The M186R mutation is the only one that occurs in the first hairpin and may disturb its β-sheet secondary structure. C203S and C206F substitute non-chelating residues for the third and fourth zinc ligands. The A200T and A200V mutations introduce bulkier sidechains that may lead to steric clashes between the two hairpins in the structure, particularly between the A200 substitutions and I196 from strand β1.

The P201H and P201S mutations may interfere with the ability of P201 to form a turn that orients the second hairpin at a right angle to the first. Two mutations R198Q and R211W have no obvious structural roles, and as such may replace residues important for molecular recognition functions of ZC4H2. Of these two sites, R211 is at the overlapping junction between the ZnF and the subsequent NLS used to direct the protein to the cell nucleus [18, 21]. A tenth mutation R213W, also in this region interferes with nuclear transport [4, 18].

### 4.2 Hierarchical folding of the ZnF domain in ZC4H2

Our results show that a 20-residue segment, ZS, is sufficient to form a high-affinity zinc-binding site (Fig. 3), but a longer 27-residue segment, ZL, is needed to obtain specific tertiary sidechain packing interactions (Fig. 2), suggesting the motif folds hierarchically. When the ZL fragment is thermally unfolded near physiological temperature (Fig. S3), the resulting molten globule may still have zinc bound. There is literature precedent for sequential non-two-state folding of ZnFs. Chelation of zinc by CCHH-type ZnFs is thought to occur in a stepwise manner with the cysteines that form the strongest interactions with the metal binding first, followed by the N-terminal His and finally the C-terminal His. A hydrophobic collapse occurs in an initial step, followed by the formation of the cysteine containing β-hairpin component of the ββα fold, and finally stabilization of the α-helix that contains the histidines [69, 70]. Similarly, there is evidence for biding of zinc by a CCCC tetradentate coordination site in the mitochondrial protein TIM10 that first involves the formation of an unstructured encounter complex as monitored by fluorimetry, followed by the development of folded structure at higher zinc concentrations in the final complex as monitored by CD [43]. Several zinc fingers show increased protein dynamics in mutant forms giving NMR spectra that approach those of unfolded proteins but retain the ability to bind zinc and adopt an overall folded structure [36, 71, 72].

To examine why the ZS fragment gives NMR spectra typical of unfolded proteins but retains CD spectra consistent with folded secondary structure (Fig. 3A), we did a simulation in which we calculated NMR structures keeping only dihedral restraints and restraints to zinc.

There is some evidence that backbone dihedral angles could be dictated by short-range interactions between residues separated by only a few positions in the polypeptide sequence [73]. When all NOE distance restraints were removed, to simulate a molten globule conformation [74] in which secondary structure is retained but sidechain interactions in the non-polar core of the protein become disordered, the resulting NMR ensemble shows a considerable spread of conformations giving a backbone of RMSD of 3.9 Å for the bundle. Nevertheless, the overall N-to C-term topology of the chain is retained (Fig. 6D) albeit with the loss of the N-terminal β-hairpin structure. The result could explain why the ZL and ZS fragments share similar CD spectra characteristic of polypeptides with folded secondary structure when zinc is bound, even though the NMR spectra suggest only ZL has specific tertiary structure. In a further simulation we removed all dihedral and distance restraints and kept only restraints between the four ligating cysteines and zinc. Even in this case, the N-to C-terminal topology of the chain is retained, although the spread in conformations is larger with an RMSD of 4.7 Å (Fig. 6E). The simulation in Fig 6E is probably unrealistic since a substantial number of residues move into disallowed regions of Ramachandran space. Taken together, these observations highlight that zinc fingers can fold sequentially, with zinc coordination forming a nidus that provides a platform for more extensive structure development.

That ZnFs fold in a hierarchical manner has implications for assessing the functions of these modules. Our approach for assessing the viability of ZnFs is to make synthetic peptide fragments corresponding to the domains, to see if these give NMR spectra typical of folded structures in the presence of zinc. Some novel ZnFs were characterized in this way, and we were able to determine that several ZnFs annotated as degenerate by UniProt were in fact functional [34–37]. The results with the ZS fragment, however, show that in some cases folding upon zinc binding could be missed by NMR which is more sensitive to fixed tertiary structure than CD which has better sensitivity to changes in secondary structure. The Protein Structure Initiative used up most resources in structural biology during the period 2000-2015 with the stated goal of cataloguing every type of protein structure [75]. Part of this effort involved the development of automation protocols to identify the most suitable candidates for structural studies. It is interesting to consider how many interesting ZnFs and other proteins could have been missed by these Big Science approaches because protein dynamics or other factors conspired to thwart the ^1^H-^15^N HSQC or crystal quality standards of that time.

### 4.3 The ZnF of ZC4H2 represents a unique sequence family amongst structural homologs

The C4 coordination site sequence pattern we established for ZC4H2 is also found in the RANBP2 family of ZnFs (Fig. 1C). Additionally, RANBP2 ZnFs have a related structure consisting of two orthogonal β-hairpins [54, 55, 76]. The prototypical member of this ZnF family, RANBP2, is a large nuclear pore protein with multiple zinc fingers that bind to the small GTPase protein Ran, which regulates the protein’s nuclear import functions [77, 78]. There are hundreds of members of the RANBP2 ZnF family [79], but their functions fall into three major categories: the ubiquitin binding NZF subfamily [76], nuclear pore proteins [78, 80], and the RNA-binding RANBP2 family [54].

The RANBP2-like family has the sequence signature W-X-C-X(2,4)-C-X3-N-X6-C-X2-C [79]. The ZnF in ZC4H2 shares this cysteine spacing but lacks the conserved tryptophan and asparagine at positions 2 and 11 of this pattern (Fig. 1C). We used the MOTIF2 server (https://www.genome.jp/tools/motif/MOTIF2.html) to search for alternative homology relationships for the ZnF of ZC4H2. If we constrained the search to include the two histidines in addition to the four cysteines with their appropriate spacing: C-X2-C-H-X3-H-X5-C-X2-C, the only hit is the ZC4H2 family. If we included the cysteines and the two unusual prolines: C-x(2)-C-x(8)-P-x-C-P-x-C, in addition to ZC4H2 we obtained hits to three RING finger families (LON peptidase 1, LON peptidase 2, and deltex E3 ubiquitin ligase 3L). While it is possible that the ZC4H2-ZnF might be evolutionarily derived from a RING finger, the homology is unlikely to be functionally relevant. This is because the RING finger motif has 8 coordinating residues that bind to two zinc ions [81]. In all three cases, the hits were to the second half of the RING finger sequence, where the first two cysteines bind to one zinc ion and the last two to the other [81]. Motif searches incorporating the conserved cysteines and residue types other than the prolines failed to find ZnF families other than ZC4H2 and the closely related homolog VAB-23. Thus, the ZnF domains from ZC4H2 and VAB-23 appear to have unique sequence properties amongst ZnFs, with the RANBP2 family amongst the closest matches in terms of the structure. It is worth noting that in addition to a distinct sequence, the structure of ZC4H2-ZnF differs from RANPB2 ZnFs by ∼4 Å RMSD and has a non-conserved C-terminal α-helix.

The best characterized ZnFs are CCHH-types that occur as DNA-binding modules in transcription factors. Yet on their own, these modules lack the ability to recognize specific dsDNA sequences, which typically requires at least two to three ZnF modules [82]. While C4 ZNFs are known to bind DNA, examples such as nuclear receptors involve a complicated sequence pattern that binds two zinc atoms and forms an α-helical structure [83]. It has been reported that RANBP2 ZnFs in nuclear pore proteins such as RANBP2 and NU153 bind to DNA but the fragments studied involved eight and four ZnF modules, respectively. There was no information provided on DNA sequence specificity, or binding affinities [78, 80]. Up until recently there were no known cases of a single C4 ZnF that binds DNA, but two examples were found in *Myobacterium smegmatis* [84]. These were 179 and 216 residue proteins, however, not the 27-residue domain found in ZC4H2. Unless it represents an unprecedented prototype, we feel it is unlikely the ZnF in ZC4H2 has a DNA-binding function since it occurs in a single copy.

A second branch of the RANBP2 family termed NZF, functions to bind ubiquitin in various post-translational modification pathways [76]. We found the possibility that ZC4H2-ZnF could bind ubiquitin intriguing in light of the literature that ZC4H2 interacts with the E3 ubiquitin ligases RNF220 [14–16] and RLIM [17] to control several pathways involved in neural development [14]. Moreover ubiquitin-binding NZF domains usually occur at the N- and C-termini of proteins like the ZNF in ZC4H2 [76]. We tested if the ZL fragment interacts with ^15^N-labeled human ubiquitin but detected no binding (Fig. S5). In retrospect, this is not surprising, since compared to the NZF sequence family, the ZnF of ZC4H2 lacks the key residues T8 and F9 (Fig. 1C) that are critical for ubiquitin binding [55, 76].

A final class of RANBP2-type ZnFs have RNA-binding functions. Using a SELEX approach it was shown that several RANBP2-type ZnFs including those from the proteins ZRANB2, EWS, RBP56, RBM5, and TEX13A, form a 1:1 complex with 6-17 nucleotide fragments of ssRNA containing the consensus sequence AGGUAA [54, 85]. A RANBP2-type ZnF in the protein FUS, recognizes the related consensus sequence NGGU [67]. Of these proteins, the testis-expressed protein TEX13A is particularly intriguing since it has the same domain organization as ZC4H2 consisting of a long coiled-coil flowed by a ZnF. However, an examination of the residues in the RANBP2 ZnFs that form key structural interactions with RNA bases [54] shows these are not conserved in ZC4H2 (cyan in Fig. 1C). Thus, while it is possible that ZC4H2 has an RNA-binding function, the nucleotide sequence of the putative RNA must be different than those bound by RANBP2 ZnFs.

Perhaps the strongest clues about the function of the ZnF in ZC4H2 come from its closest homolog, the *C. elegans* protein Vab-23. Homology for the two proteins is particularly high for the ZnF and subsequent NLS regions (Fig. 1C), which were originally used to infer the presence of a ZnF motif in ZC4H2 [21]. Vab-23 is a regulator of epidermal morphogenesis in *C. elegans* embryos and its ZnF domain is required for this function. The protein is localized to nuclear speckles in a manner consistent with pre-mRNA processing or mRNA regulation [21] suggesting it binds RNA. Thus, an RNA-binding role for the ZnF of ZC4H2 is plausible albeit with RNA sequence determinants different from those of the RANBP2 domains. Alternatively, since ZC4H2 has both unique sequence and structural properties it may have an alternative function.

## 5. CONCLUSIONS

In this work we showed that ZC4H2 contains a genuine ZnF which has implications for the structure and function of the protein, as well as its role in disease. Establishing the presence of a genuine ZnF in the ZC4H2 protein was not trivial, since the AI-based AF3 program predicts folded structures for peptides of similar lengths with random sequences that are unlikely to be folded. Even using experimental methods, establishing the presence of a functional ZnF proved difficult since the UniProt-specified domain boundaries give a peptide that binds zinc but forms a molten globule, lacking specific sidechain packing until the fragment is appended at its N- and C-terminal ends. Thus, testing the viability of a ZnF can require multiple experimental techniques as well as optimization of domain boundaries to achieve proper folding. The presence of a ZnF in ZC4H2 raises the possibility that this domain could participate in its protein-protein interactions, including with SMAD proteins [9], the TRPV4 ion channel [11], and the RNF220 and RLIM E3 ubiquitin ligases [17], amongst others.

Nine ZARD mutations occur in the ZnF and while most of these are likely to perturb the structure, at least one mutation, R198Q, occurs at a position that does not appear to serve a structural role. As such the mutation could disrupt molecular interactions involving the ZnF. Two other mutations R211W and R213W near the C-terminus also do not appear to have structural roles but overlap with a NLS sequence [18] and could affect binding interactions or nuclear import of ZC4H2.

The sequence and NMR structure of the ZC4H2 ZnF is similar those from the RANBP2 family but has important differences in sequence and structure. Because the ZnF modules share a similar sequence spacing of the four Zn^2+^-binding cysteine ligands, we looked for functional commonalities. Unlike the NZF branch of the RANBP2 family the ZC4H2 ZnF does not bind ubiquitin. The ZC4H2 ZnF is unlikely to bind DNA since it occurs as a single copy and is probably too short to provide an epitope for specific DNA-binding [82, 84]. It is possible that ZC4H2 ZnF binds RNA since it has a high proportion of positively charged residues and a fold used for this function in other ZnFs, but the RNA sequence-specificity must be different than that for the RNA-binding branch of the RANBP2 family since key binding residues are missing. The current structural work sets a foundation for determining the function of the ZnF in the ZC4H2 protein.

## ACKNOWLEDGMENTS

We thank Prof. Irina Bezsanova and Daniel Fairchild for a sample of ^15^N-ubiquitin, and Prof. Carolyn Teschke for use of her CD and UV-Vis instruments.

## AUTHOR CONTRIBUTIONS

Conceptualization: A.T.A and R.E.H.; discovery: R.E.H.; methodology: A.T.A. and A.J.R; investigations: A.T.A, R.E.H. and A.J.R.; formal analysis: A.T.A, R.E.H. and A.J.R.; resources: A.T.A; writing – original draft preparation: A.T.A; writing – reviewing and editing, A.T.A, R.E.H. and A.J.R.. All authors have read and agreed to the published version of the manuscript.

## SUPPLEMENTARY DATA

Five figures and one table are available at http://www.xxx

## References

1. Piccolo, G.; d’Annunzio, G.; Amadori, E.; Riva, A.; Borgia, P.; Tortora, D.; Maghnie, M.; Minetti, C.; Gitto, E.; Iacomino, M.; Baldassari, S.; Fiorillo, C.; Zara, F.; Striano, P.; Salpietro, V., Neuromuscular and Neuroendocrinological Features Associated With ZC4H2-Related Arthrogryposis Multiplex Congenita in a Sicilian Family: A Case Report. Front Neurol 2021, 12, 704747.

2. Peters, S.; Sportiello, K.; Mandalapu, S.; Nguyen, A.; Carrier, R.; Dickinson, C.; Paciorkowski, A.; Bearden, D., Genotype-Phenotype Correlations and Sex Differences in ZC4H2-Associated Rare Disorder. Pediatr Neurol 2024, 158, 100–112.

3. Frints, S. G. M.; Hennig, F.; Colombo, R.; Jacquemont, S.; Terhal, P.; Zimmerman, H. H.; Hunt, D.; Mendelsohn, B. A.; Kordass, U.; Webster, R.; Sinnema, M.; Abdul-Rahman, O.; Suckow, V.; Fernandez-Jaen, A.; van Roozendaal, K.; Stevens, S. J. C.; Macville, M. V. E.; Al-Nasiry, S.; van Gassen, K.; Utzig, N.; Koudijs, S. M.; McGregor, L.; Maas, S. M.; Baralle, D.; Dixit, A.; Wieacker, P.; Lee, M.; Lee, A. S.; Engle, E. C.; Houge, G.; Gradek, G. A.; Douglas, A. G. L.; Longman, C.; Joss, S.; Velasco, D.; Hennekam, R. C.; Hirata, H.; Kalscheuer, V. M., Deleterious de novo variants of X-linked ZC4H2 in females cause a variable phenotype with neurogenic arthrogryposis multiplex congenita. Hum Mutat 2019, 40, (12), 2270–2285.

4. Hirata, H.; Nanda, I.; van Riesen, A.; McMichael, G.; Hu, H.; Hambrock, M.; Papon, M. A.; Fischer, U.; Marouillat, S.; Ding, C.; Alirol, S.; Bienek, M.; Preisler-Adams, S.; Grimme, A.; Seelow, D.; Webster, R.; Haan, E.; MacLennan, A.; Stenzel, W.; Yap, T. Y.; Gardner, A.; Nguyen, L. S.; Shaw, M.; Lebrun, N.; Haas, S. A.; Kress, W.; Haaf, T.; Schellenberger, E.; Chelly, J.; Viot, G.; Shaffer, L. G.; Rosenfeld, J. A.; Kramer, N.; Falk, R.; El-Khechen, D.; Escobar, L. F.; Hennekam, R.; Wieacker, P.; Hubner, C.; Ropers, H. H.; Gecz, J.; Schuelke, M.; Laumonnier, F.; Kalscheuer, V. M., ZC4H2 mutations are associated with arthrogryposis multiplex congenita and intellectual disability through impairment of central and peripheral synaptic plasticity. Am J Hum Genet 2013, 92, (5), 681–95.

5. Wieacker, P.; Wolff, G.; Wienker, T. F.; Sauer, M., A new X-linked syndrome with muscle atrophy, congenital contractures, and oculomotor apraxia. Am J Med Genet 1985, 20, (4), 597–606.

6. Ibarra-Ramirez, M.; Fernandez-de-Luna, M. L.; Campos-Acevedo, L. D.; Arenas-Estala, J.; Martinez-de-Villarreal, L. E.; Rodriguez-Garza, C.; DeLagarza-Pineda, O.; Mohamed-Noriega, J., Optic nerve abnormalities in female-restricted Wieacker-Wolff syndrome by a novel variant in the ZC4H2 gene. Ophthalmic Genet 2023, 44, (5), 465–468.

7. Wakabayashi, T.; Mizukami, M.; Terada, K.; Ishikawa, A.; Hinotsu, S.; Kobayashi, M.; Kato, K.; Ogi, T.; Tsugawa, T.; Sakurai, A., A novel ZC4H2 variant in a female with severe respiratory complications. Brain Dev 2022, 44, (8), 571–577.

8. Del Toro, N.; Shrivastava, A.; Ragueneau, E.; Meldal, B.; Combe, C.; Barrera, E.; Perfetto, L.; How, K.; Ratan, P.; Shirodkar, G.; Lu, O.; Meszaros, B.; Watkins, X.; Pundir, S.; Licata, L.; Iannuccelli, M.; Pellegrini, M.; Martin, M. J.; Panni, S.; Duesbury, M.; Vallet, S. D.; Rappsilber, J.; Ricard-Blum, S.; Cesareni, G.; Salwinski, L.; Orchard, S.; Porras, P.; Panneerselvam, K.; Hermjakob, H., The IntAct database: efficient access to fine-grained molecular interaction data. Nucleic Acids Res 2022, 50, (D1), D648–D653.

9. Ma, P.; Ren, B.; Yang, X.; Sun, B.; Liu, X.; Kong, Q.; Li, C.; Mao, B., ZC4H2 stabilizes Smads to enhance BMP signalling, which is involved in neural development in Xenopus. Open Biol 2017, 7, (8).

10. Kobayashi, S.; Sato, A.; Chiba, Y.; Adachi, N.; Kakimoto, Y.; Suzuki, H.; Yamada, M.; Kosaki, K.; Tanaka, H., Wieacker-Wolff syndrome with hyperinsulinemic hypoglycemia successfully treated using diazoxide: A case report. Clin Pediatr Endocrinol 2025, 34, (1), 70–76.

11. Vangeel, L.; Janssens, A.; Lemmens, I.; Lievens, S.; Tavernier, J.; Voets, T., The Zinc-Finger Domain Containing Protein ZC4H2 Interacts with TRPV4, Enhancing Channel Activity and Turnover at the Plasma Membrane. Int J Mol Sci 2020, 21, (10).

12. Cho, T. J.; Matsumoto, K.; Fano, V.; Dai, J.; Kim, O. H.; Chae, J. H.; Yoo, W. J.; Tanaka, Y.; Matsui, Y.; Takigami, I.; Monges, S.; Zabel, B.; Shimizu, K.; Nishimura, G.; Lausch, E.; Ikegawa, S., TRPV4-pathy manifesting both skeletal dysplasia and peripheral neuropathy: a report of three patients. Am J Med Genet A 2012, 158A, (4), 795–802.

13. Briscoe, J.; Therond, P. P., The mechanisms of Hedgehog signalling and its roles in development and disease. Nat Rev Mol Cell Biol 2013, 14, (7), 416–29.

14. Kim, J.; Choi, T. I.; Park, S.; Kim, M. H.; Kim, C. H.; Lee, S., Rnf220 cooperates with Zc4h2 to specify spinal progenitor domains. Development 2018, 145, (17).

15. Ma, P.; Mao, B., The many faces of the E3 ubiquitin ligase, RNF220, in neural development and beyond. Dev Growth Differ 2022, 64, (2), 98-105.

16. Ma, P.; Song, N. N.; Cheng, X.; Zhu, L.; Zhang, Q.; Zhang, L. L.; Yang, X.; Wang, H.; Kong, Q.; Shi, D.; Ding, Y. Q.; Mao, B., ZC4H2 stabilizes RNF220 to pattern ventral spinal cord through modulating Shh/Gli signaling. J Mol Cell Biol 2020, 12, (5), 337–344.

17. Li, Y.; Yang, C.; Wang, H.; Zhao, L.; Kong, Q.; Cang, Y.; Zhao, S.; Lv, L.; Li, Y.; Mao, B.; Ma, P., Sequential stabilization of RNF220 by RLIM and ZC4H2 during cerebellum development and Shh-group medulloblastoma progression. J Mol Cell Biol 2022, 14, (1).

18. Wang, D.; Hu, D.; Guo, Z.; Hu, R.; Wang, Q.; Liu, Y.; Liu, M.; Meng, Z.; Yang, H.; Zhang, Y.; Cai, F.; Zhou, W.; Song, W., A novel de novo nonsense mutation in ZC4H2 causes Wieacker-Wolff Syndrome. Mol Genet Genomic Med 2020, 8, (2), e1100.

19. Hirosawa, M.; Nagase, T.; Ishikawa, K.; Kikuno, R.; Nomura, N.; Ohara, O., Characterization of cDNA clones selected by the GeneMark analysis from size-fractionated cDNA libraries from human brain. DNA Res 1999, 6, (5), 329–36.

20. Wang, Y.; Han, K. J.; Pang, X. W.; Vaughan, H. A.; Qu, W.; Dong, X. Y.; Peng, J. R.; Zhao, H. T.; Rui, J. A.; Leng, X. S.; Cebon, J.; Burgess, A. W.; Chen, W. F., Large scale identification of human hepatocellular carcinoma-associated antigens by autoantibodies. J Immunol 2002, 169, (2), 1102–9.

21. Pellegrino, M. W.; Gasser, R. B.; Sprenger, F.; Stetak, A.; Hajnal, A., The conserved zinc finger protein VAB-23 is an essential regulator of epidermal morphogenesis in Caenorhabditis elegans. Dev Biol 2009, 336, (1), 84–93.

22. Berg, J. M., Proposed structure for the zinc-binding domains from transcription factor IIIA and related proteins. Proc Natl Acad Sci U S A 1988, 85, (1), 99–102.

23. Klug, A., The discovery of zinc fingers and their applications in gene regulation and genome manipulation. Annu Rev Biochem 2010, 79, 213–31.

24. Kluska, K.; Adamczyk, J.; Krezel, A., Metal binding properties, stability and reactivity of zinc fingers. Coordination Chemistry Reviews 2018, 367, 18–64.

25. Lee, M. S.; Gippert, G. P.; Soman, K. V.; Case, D. A.; Wright, P. E., Three-dimensional solution structure of a single zinc finger DNA-binding domain. Science 1989, 245, (4918), 635–7.

26. Michalek, J. L.; Besold, A. N.; Michel, S. L., Cysteine and histidine shuffling: mixing and matching cysteine and histidine residues in zinc finger proteins to afford different folds and function. Dalton Trans 2011, 40, (47), 12619–32.

27. Neuhaus, D., Zinc finger structure determination by NMR: Why zinc fingers can be a handful. Prog Nucl Magn Reson Spectrosc 2022, 130–131, 62-105.

28. Abbehausen, C., Zinc finger domains as therapeutic targets for metal-based compounds - an update. Metallomics 2019, 11, (1), 15–28.

29. Padjasek, M.; Kocyla, A.; Kluska, K.; Kerber, O.; Tran, J. B.; Krezel, A., Structural zinc binding sites shaped for greater works: Structure-function relations in classical zinc finger, hook and clasp domains. J Inorg Biochem 2020, 204, 110955.

30. Krishna, S. S.; Majumdar, I.; Grishin, N. V., Structural classification of zinc fingers: survey and summary. Nucleic Acids Res 2003, 31, (2), 532–50.

31. Lambert, S. A.; Jolma, A.; Campitelli, L. F.; Das, P. K.; Yin, Y.; Albu, M.; Chen, X.; Taipale, J.; Hughes, T. R.; Weirauch, M. T., The Human Transcription Factors. Cell 2018, 172, (4), 650–665.

32. Kluska, K.; Adamczyk, J.; Krezel, A., Metal binding properties of zinc fingers with a naturally altered metal binding site. Metallomics 2018, 10, (2), 248–263.

33. Callaway, E., Revolutionary cryo-EM is taking over structural biology. Nature 2020, 578, (7794), 201.

34. Harris, R. E.; Whitehead, R. D., 3rd; Alexandrescu, A. T., Solution structure of the Z0 domain from transcription repressor BCL11A sheds light on the sequence properties of protein-binding zinc fingers. Protein Sci 2025, 34, (4), e70097.

35. Matousek, W. M.; Alexandrescu, A. T., NMR structure of the C-terminal domain of SecA in the free state. Biochim Biophys Acta 2004, 1702, (2), 163–71.

36. Rua, A. J.; Alexandrescu, A. T., Formerly degenerate seventh zinc finger domain from transcription factor ZNF711 rehabilitated by experimental NMR structure. Protein Sci 2024, 33, (9), e5149.

37. Rua, A. J.; Whitehead, R. D., 3rd; Alexandrescu, A. T., NMR structure verifies the eponymous zinc finger domain of transcription factor ZNF750. J Struct Biol X 2023, 8, 100093.

38. Walker, J. M., The Bicinchoninic Acid (BCA) Assay for Protein Quantitation. In The Protein Protocols Handbook, 2nd ed.; Walker, J. M., Ed. Humana Press: Totowa, NJ, 2002; pp 11-15.

39. Wishart, D. S.; Bigam, C. G.; Yao, J.; Abildgaard, F.; Dyson, H. J.; Oldfield, E.; Markley, J. L.; Sykes, B. D., 1H, 13C and 15N chemical shift referencing in biomolecular NMR. J Biomol NMR 1995, 6, (2), 135-40.

40. Harprecht, C.; Okifo, O.; Robbins, K. J.; Motwani, T.; Alexandrescu, A. T.; Teschke, C. M., Contextual Role of a Salt Bridge in the Phage P22 Coat Protein I-Domain. J Biol Chem 2016, 291, (21), 11359–72.

41. Whitehead, R. D., 3rd; Teschke, C. M.; Alexandrescu, A. T., Pulse-field gradient nuclear magnetic resonance of protein translational diffusion from native to non-native states. Protein Sci 2022, 31, (5), e4321.

42. Vranken, W. F.; Boucher, W.; Stevens, T. J.; Fogh, R. H.; Pajon, A.; Llinas, M.; Ulrich, E. L.; Markley, J. L.; Ionides, J.; Laue, E. D., The CCPN data model for NMR spectroscopy: development of a software pipeline. Proteins 2005, 59, (4), 687–96.

43. Ivanova, E.; Ball, M.; Lu, H., Zinc binding of Tim10: Evidence for existence of an unstructured binding intermediate for a zinc finger protein. 2008, 71, (1), 467-475.

44. Abramson, J.; Adler, J.; Dunger, J.; Evans, R.; Green, T.; Pritzel, A.; Ronneberger, O.; Willmore, L.; Ballard, A. J.; Bambrick, J.; Bodenstein, S. W.; Evans, D. A.; Hung, C. C.; O’Neill, M.; Reiman, D.; Tunyasuvunakool, K.; Wu, Z.; Zemgulyte, A.; Arvaniti, E.; Beattie, C.; Bertolli, O.; Bridgland, A.; Cherepanov, A.; Congreve, M.; Cowen-Rivers, A. I.; Cowie, A.; Figurnov, M.; Fuchs, F. B.; Gladman, H.; Jain, R.; Khan, Y. A.; Low, C. M. R.; Perlin, K.; Potapenko, A.; Savy, P.; Singh, S.; Stecula, A.; Thillaisundaram, A.; Tong, C.; Yakneen, S.; Zhong, E. D.; Zielinski, M.; Zidek, A.; Bapst, V.; Kohli, P.; Jaderberg, M.; Hassabis, D.; Jumper, J. M., Accurate structure prediction of biomolecular interactions with AlphaFold 3. Nature 2024, 630, (8016), 493–500.

45. Shen, Y.; Bax, A., Protein backbone and sidechain torsion angles predicted from NMR chemical shifts using artificial neural networks. J Biomol NMR 2013, 56, (3), 227–41.

46. Wuthrich, K.; Billeter, M.; Braun, W., Pseudo-structures for the 20 common amino acids for use in studies of protein conformations by measurements of intramolecular proton-proton distance constraints with nuclear magnetic resonance. J Mol Biol 1983, 169, (4), 949–61.

47. Ramboarina, S.; Druillennec, S.; Morellet, N.; Bouaziz, S.; Roques, B. P., Target specificity of human immunodeficiency virus type 1 NCp7 requires an intact conformation of its CCHC N-terminal zinc finger. J Virol 2004, 78, (12), 6682–7.

48. Tejero, R.; Snyder, D.; Mao, B.; Aramini, J. M.; Montelione, G. T., PDBStat: a universal restraint converter and restraint analysis software package for protein NMR. J Biomol NMR 2013, 56, (4), 337–51.

49. Maciejewski, M. W.; Schuyler, A. D.; Gryk, M. R.; Moraru, II; Romero, P. R.; Ulrich, E. L.; Eghbalnia, H. R.; Livny, M.; Delaglio, F.; Hoch, J. C., NMRbox: A Resource for Biomolecular NMR Computation. Biophys J 2017, 112, (8), 1529–1534.

50. Bermejo, G. A.; Tjandra, N.; Clore, G. M.; Schwieters, C. D., Xplor-NIH: Better parameters and protocols for NMR protein structure determination. Protein Sci 2024, 33, (4), e4922.

51. Rieping, W.; Habeck, M.; Bardiaux, B.; Bernard, A.; Malliavin, T. E.; Nilges, M., ARIA2: automated NOE assignment and data integration in NMR structure calculation. Bioinformatics 2007, 23, (3), 381–2.

52. Whitehead, R. D., 3rd; Teschke, C. M.; Alexandrescu, A. T., NMR Mapping of Disordered Segments from a Viral Scaffolding Protein Enclosed in a 23 MDa Procapsid. Biophys J 2019, 117, (8), 1387-1392.

53. Alexandrescu, A. T.; Evans, P. A.; Pitkeathly, M.; Baum, J.; Dobson, C. M., Structure and dynamics of the acid-denatured molten globule state of alpha-lactalbumin: a two-dimensional NMR study. Biochemistry 1993, 32, (7), 1707–18.

54. Nguyen, C. D.; Mansfield, R. E.; Leung, W.; Vaz, P. M.; Loughlin, F. E.; Grant, R. P.; Mackay, J. P., Characterization of a family of RanBP2-type zinc fingers that can recognize single-stranded RNA. J Mol Biol 2011, 407, (2), 273–83.

55. Wang, B.; Alam, S. L.; Meyer, H. H.; Payne, M.; Stemmler, T. L.; Davis, D. R.; Sundquist, W. I., Structure and ubiquitin interactions of the conserved zinc finger domain of Npl4. J Biol Chem 2003, 278, (22), 20225–34.

56. Curtis, D.; Treiber, D. K.; Tao, F.; Zamore, P. D.; Williamson, J. R.; Lehmann, R., A CCHC metal-binding domain in Nanos is essential for translational regulation. EMBO J 1997, 16, (4), 834–43.

57. Croke, R. L.; Patil, S. M.; Quevreaux, J.; Kendall, D. A.; Alexandrescu, A. T., NMR determination of pKa values in alpha-synuclein. Protein Sci 2011, 20, (2), 256–69.

58. Jha, S.; Snell, J. M.; Sheftic, S. R.; Patil, S. M.; Daniels, S. B.; Kolling, F. W.; Alexandrescu, A. T., pH dependence of amylin fibrillization. Biochemistry 2014, 53, (2), 300–10.

59. Nomura, A.; Sugiura, Y., Contribution of individual zinc ligands to metal binding and peptide folding of zinc finger peptides. Inorg Chem 2002, 41, (14), 3693–8.

60. Krizek, B. A.; Merkle, D. L.; Berg, J. M., Ligand Variation and Metal Ion Binding Specificity in Zinc Finger Peptides. Inorg Chem 1993, 32, 937–940.

61. Sivo, V.; D’Abrosca, G.; Russo, L.; Iacovino, R.; Pedone, P. V.; Fattorusso, R.; Isernia, C.; Malgieri, G., Co(II) Coordination in Prokaryotic Zinc Finger Domains as Revealed by UV-Vis Spectroscopy. Bioinorg Chem Appl 2017, 2017, 1527247.

62. Gasteiger, E.; Hoogland, C.; Gattiker, A.; Duvaud, S.; Wilkins, M. R.; Appel, R. D.; Bairoch, A., Protein Identification and Analysis Tools on the Expasy Server. In The Proteomics Protocols Handbook, Walker, J. M., Ed. Humana Press: Totowa, NJ, 2005.

63. Orengo, C. A.; Michie, A. D.; Jones, S.; Jones, D. T.; Swindells, M. B.; Thornton, J. M., CATH--a hierarchic classification of protein domain structures. Structure 1997, 5, (8), 1093–108.

64. Hassel, K. R.; Brito-Estrada, O.; Makarewich, C. A., Microproteins: Overlooked regulators of physiology and disease. iScience 2023, 26, (6), 106781.

65. Pennisi, E., ’Dark proteome’ survey reveals thousands of new human genes. Science 2024, 386, (6725), 951–952.

66. Li, Y.; Okatsu, K.; Fukai, S.; Sato, Y., Structural basis for specific recognition of K6-linked polyubiquitin chains by the TAB2 NZF domain. Biophys J 2021, 120, (16), 3355–3362.

67. Loughlin, F. E.; Lukavsky, P. J.; Kazeeva, T.; Reber, S.; Hock, E. M.; Colombo, M.; Von Schroetter, C.; Pauli, P.; Clery, A.; Muhlemann, O.; Polymenidou, M.; Ruepp, M. D.; Allain, F. H., The Solution Structure of FUS Bound to RNA Reveals a Bipartite Mode of RNA Recognition with Both Sequence and Shape Specificity. Mol Cell 2019, 73, (3), 490–504 e6.

68. Baker, D.; Sali, A., Protein structure prediction and structural genomics. Science 2001, 294, (5540), 93–6.

69. Miura, T.; Satoh, T.; Takeuchi, H., Role of metal-ligand coordination in the folding pathway of zinc finger peptides. Biochim Biophys Acta 1998, 1384, (1), 171–9.

70. Li, W.; Zhang, J.; Wang, J.; Wang, W., Metal-coupled folding of Cys2His2 zinc-finger. J Am Chem Soc 2008, 130, (3), 892–900.

71. Cordier, F.; Vinolo, E.; Veron, M.; Delepierre, M.; Agou, F., Solution structure of NEMO zinc finger and impact of an anhidrotic ectodermal dysplasia with immunodeficiency-related point mutation. J Mol Biol 2008, 377, (5), 1419–32.

72. Simpson, R. J.; Cram, E. D.; Czolij, R.; Matthews, J. M.; Crossley, M.; Mackay, J. P., CCHX zinc finger derivatives retain the ability to bind Zn(II) and mediate protein-DNA interactions. J Biol Chem 2003, 278, (30), 28011–8.

73. Shortle, D., Propensities, probabilities, and the Boltzmann hypothesis. Protein Sci 2003, 12, (6), 1298–302.

74. Dolgikh, D. A.; Gilmanshin, R. I.; Brazhnikov, E. V.; Bychkova, V. E.; Semisotnov, G. V.; Venyaminov, S.; Ptitsyn, O. B., Alpha-Lactalbumin: compact state with fluctuating tertiary structure? FEBS Lett 1981, 136, (2), 311–5.

75. Petsko, G. A., An idea whose time has gone. Genome Biol 2007, 8, (6), 107.

76. Michel, M. A.; Scutts, S.; Komander, D., Secondary interactions in ubiquitin-binding domains achieve linkage or substrate specificity. Cell Rep 2024, 43, (8), 114545.

77. Yaseen, N. R.; Blobel, G., Two distinct classes of Ran-binding sites on the nucleoporin Nup-358. Proc Natl Acad Sci U S A 1999, 96, (10), 5516–21.

78. Yokoyama, N.; Hayashi, N.; Seki, T.; Pante, N.; Ohba, T.; Nishii, K.; Kuma, K.; Hayashida, T.; Miyata, T.; Aebi, U.;, et al., A giant nucleopore protein that binds Ran/TC4. Nature 1995, 376, (6536), 184–8.

79. Higa, M. M.; Alam, S. L.; Sundquist, W. I.; Ullman, K. S., Molecular characterization of the Ran-binding zinc finger domain of Nup153. J Biol Chem 2007, 282, (23), 17090–100.

80. Sukegawa, J.; Blobel, G., A nuclear pore complex protein that contains zinc finger motifs, binds DNA, and faces the nucleoplasm. Cell 1993, 72, (1), 29–38.

81. Garcia-Barcena, C.; Osinalde, N.; Ramirez, J.; Mayor, U., How to Inactivate Human Ubiquitin E3 Ligases by Mutation. Front Cell Dev Biol 2020, 8, 39.

82. Persikov, A. V.; Singh, M., De novo prediction of DNA-binding specificities for Cys2His2 zinc finger proteins. Nucleic Acids Res 2014, 42, (1), 97–108.

83. Schwabe, J. W.; Rhodes, D., Beyond zinc fingers: steroid hormone receptors have a novel structural motif for DNA recognition. Trends Biochem Sci 1991, 16, (8), 291–6.

84. Ghosh, S.; Chatterji, D., Two zinc finger proteins from Mycobacterium smegmatis: DNA binding and activation of transcription. Genes Cells 2017, 22, (8), 699–714.

85. Loughlin, F. E.; Mansfield, R. E.; Vaz, P. M.; McGrath, A. P.; Setiyaputra, S.; Gamsjaeger, R.; Chen, E. S.; Morris, B. J.; Guss, J. M.; Mackay, J. P., The zinc fingers of the SR-like protein ZRANB2 are single-stranded RNA-binding domains that recognize 5’ splice site-like sequences. Proc Natl Acad Sci U S A 2009, 106, (14), 5581–6.

86. Singh, B. B.; Patel, H. H.; Roepman, R.; Schick, D.; Ferreira, P. A., The zinc finger cluster domain of RanBP2 is a specific docking site for the nuclear export factor, exportin-1. J Biol Chem 1999, 274, (52), 37370–8.

87. Alam, S. L.; Sun, J.; Payne, M.; Welch, B. D.; Blake, B. K.; Davis, D. R.; Meyer, H. H.; Emr, S. D.; Sundquist, W. I., Ubiquitin interactions of NZF zinc fingers. EMBO J 2004, 23, (7), 1411–21.

