## Supplementary Information for "Zinc-induced folding and solution structure of the eponymous novel zinc finger from the ZC4H2 protein"

**Table of contents:**

**Figure S1 – DOSY spectrum of ZC4H2-ZL**
**Figure S2 – $^{13}$C- and $^{15}$N-HSQC spectra of ZL-$Zn^{2+}$ with assigned fingerprint regions**
**Figure S3 – Thermal unfolding of ZL-$Zn^{2+}$ by NMR**
**Figure S4 – NMR secondary structure and NOEs supporting a short C-terminal $\alpha$-helix**
**Figure S5 – NMR fails to detect interaction between ZC4H2-ZL and ubiquitin**

**Table S1 – Statistics for the 20 best NMR structures of the ZL fragment.**

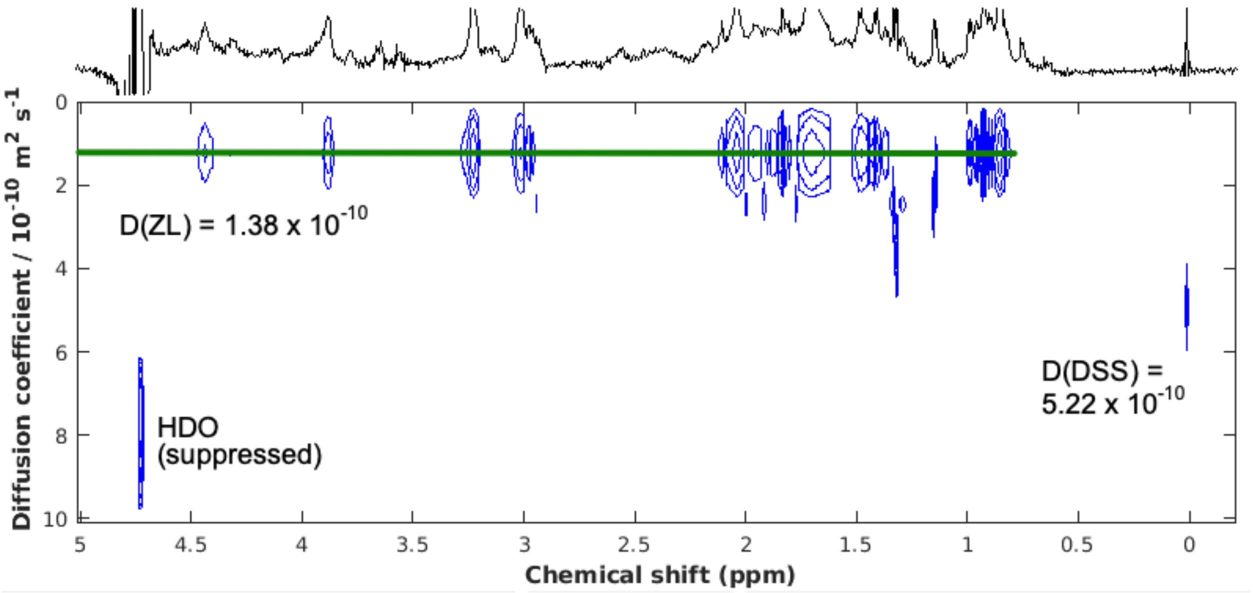


**Supplementary Figure S1 DOSY spectrum of ZC4H2-ZL.** A 0.3 mM ZL sample in $D_2O$ at pH 6.87 with equimolar $ZnCl_2$ was used for the experiments. Although most NMR work for ZL was done at a 10 C, a temperature of 25 C was used for the DOSY experiments to avoid convection artefacts. NMR diffusion data were acquired on a 600 MHz Varian Inova spectrometer with the *Doneshot* pulse sequence and processed with the GNAT program on the NMRbox platform. As indicated in the figure, diffusion coefficients $D_{DSS}$ = 5.22 x $10^{-10}$ $m^2$ $s^{-1}$ and $D_{ZL}$ = 1.38 x $10^{-10}$ $m^2$ $s^{-1}$ were obtained for the DSS standard and ZL-$Zn^{2+}$, respectively. From the $R_h$ (radius of hydration) value of 3.34 Å for DSS and the formula $R_{h, ZL} = (D_{DSS}/D_{Z0})*R_{h,DSS}$, an Rh value of 12.6 Å was calculated for the folded $Zn^{2+}$-bound ZL peptide, consistent with a monomer (an $R_h$ of 12.35 Å is expected for a 27-residue monomer, compared to 15.10 Å for a 54-residue dimer).

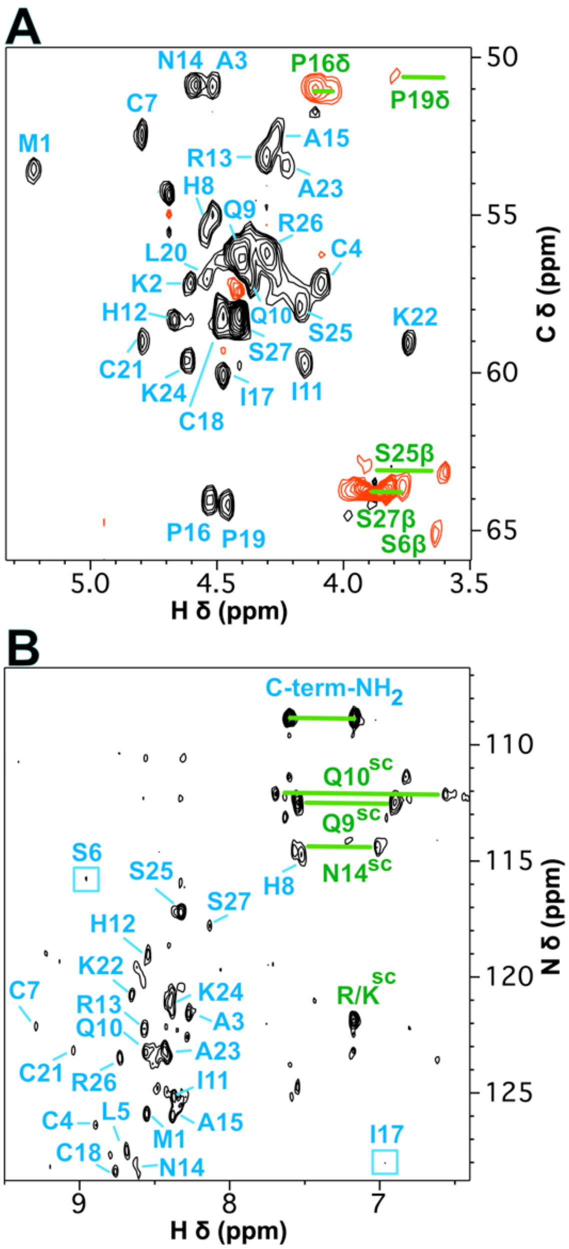


**Supplementary Figure S2 $^{13}$C- and $^{15}$N-HSQC spectra of ZL-$Zn^{2+}$ with assigned fingerprint regions.** The $^{1}$H-$^{13}$C (**A**) and $^{1}$H-$^{15}$N HSQC (**B**) spectra were collected on samples at natural isotope abundance. Backbone Hα-Cα and H-N crosspeaks are labeled with their sequence-specific NMR assignments in blue, correlations due to sidechains in green. The multiplicity-edited $^{1}$H-$^{13}$C HSQC in (A) was recorded on a Bruker 600 MHz Neo spectrometer equipped with a cryoprobe on a 0.9 mM ZL sample with equimolar $ZnCl_2$ dissolved in $D_2O$. In the multiplicity-edited $^{1}$H-$^{13}$C HSQC spectrum, CH and $CH_3$ groups give black contours, whereas $CH_2$ groups red contours The $^{1}$H-$^{15}$N HSQC in (B) was recorded on a Bruker 800 MHz Neo spectrometer with a cryoprobe on a 1.6 mM ZL sample dissolved in $H_2O$, at pH 5.9 and a temperature of 10 C in a total acquisition time of 12 h. Because of the low sensitivity of the $^{1}$H-$^{15}$N HSQC at natural abundance, assignments of weak peaks in the spectrum should be considered tentative.

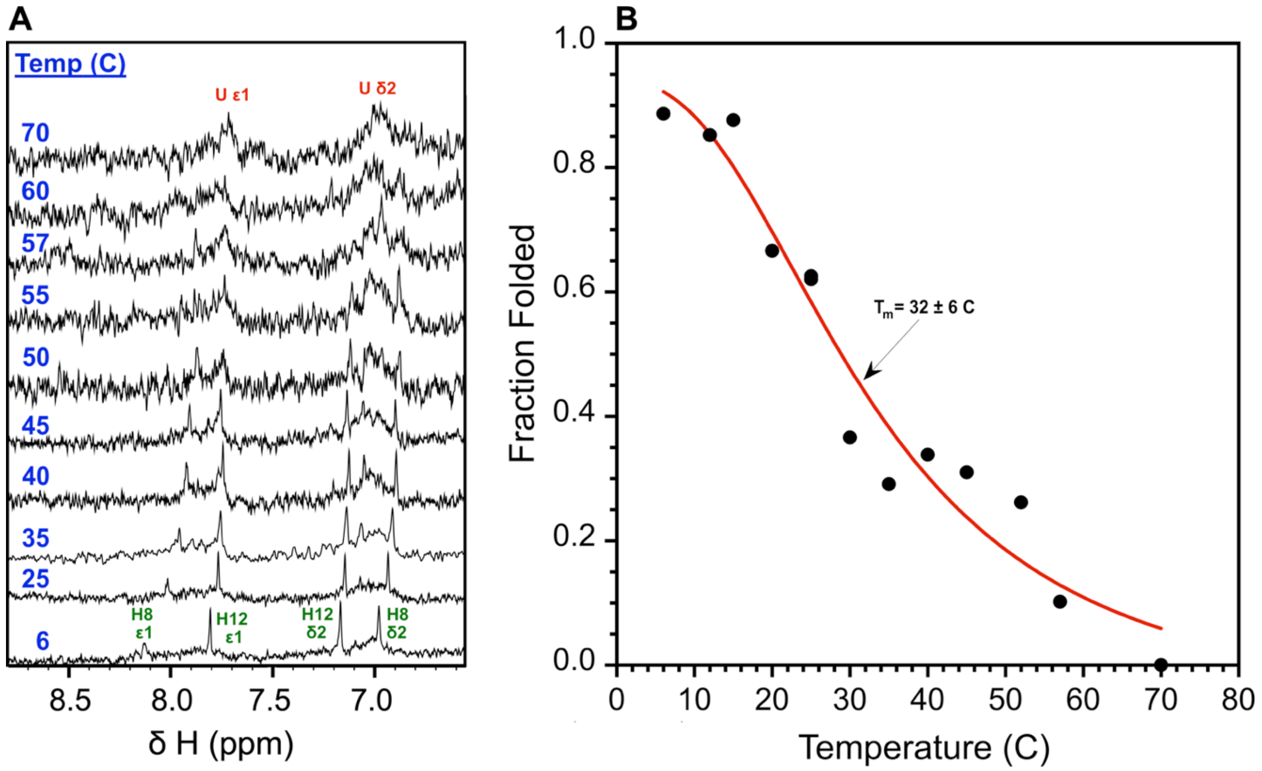


**Supplementary Figure S3 Thermal unfolding of ZL-Zn$^{2+}$ by NMR** (**A**) Thermal melt monitored by 1d $^1$H-NMR spectra of the aromatic resonances from the two histidines in ZL-Zn$^{2+}$. The sample was 300 μM ZL-Zn$^{2+}$ at pH 6.9. Each of the two histidines gives resolved Hε1 and Hδ2 signals in the folded state (green). With increasing temperature, the Hε1 signal from the unfolded state of both histidines (red, Uε1) overlaps with the folded Hε1 signal for H12. The unresolved signal from the Hδ2 resonances of the two histidines in the unfolded state (red, Uδ2) is positioned between the two folded-state Hδ2 resonances and was used to calculate the fraction of unfolded protein at each temperature using the integration routine of the program iNMR. (**B**) Fitting of the fraction of folded protein versus temperature to a sigmoidal curve gave a midpoint for thermal unfolding of 32 ± 6 C.

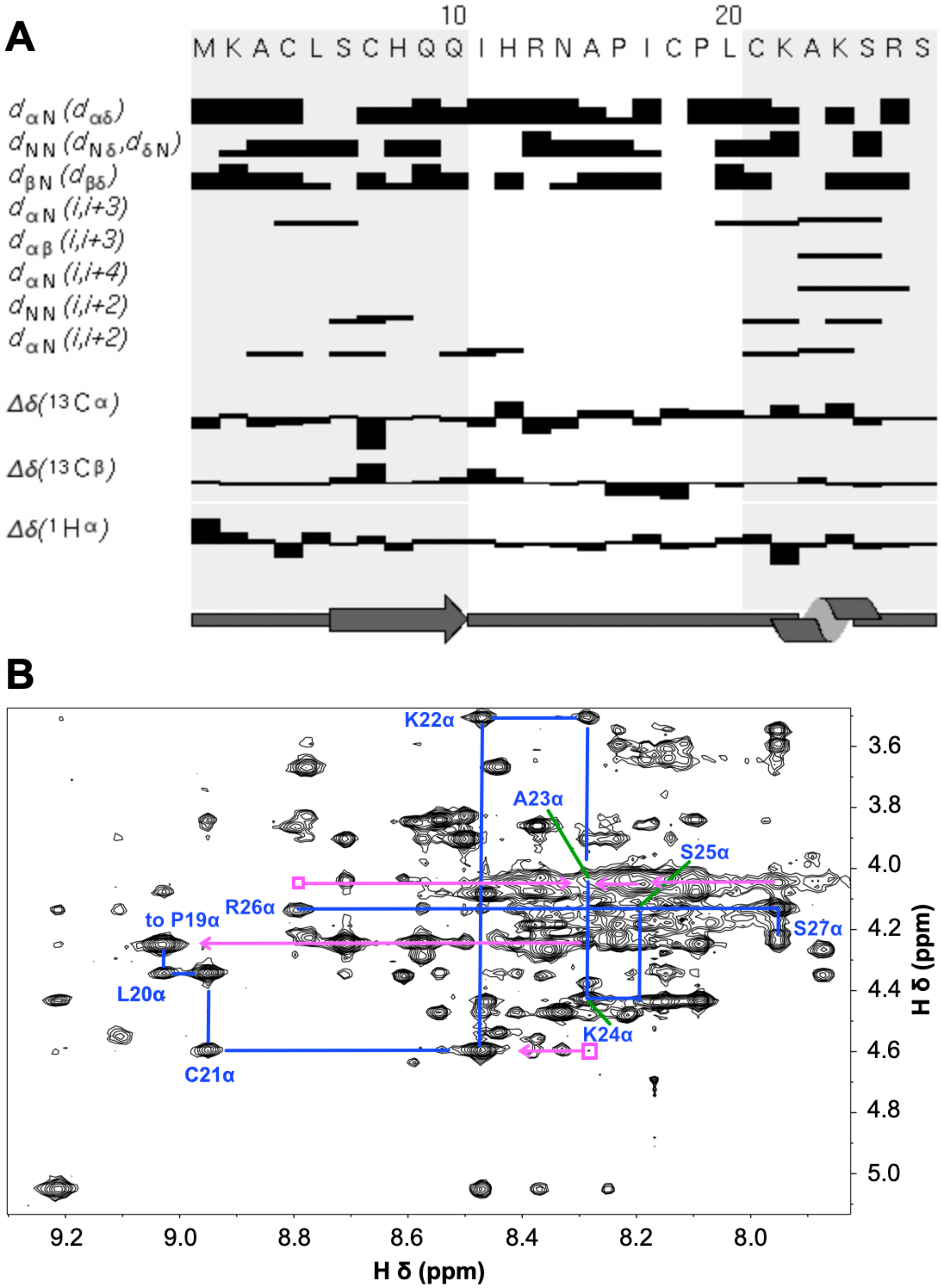


**Supplementary Figure S4 NMR secondary structure and NOEs supporting a short C-terminal α-helix** (**A**) Wüthrich diagram showing short-range NOEs, chemical shift indices, and consensus NMR-derived secondary structure. (**B**) NOEs supporting a short one-turn α-helix at the C-terminus. Labels indicate intraresidue HN-Hα NOEs, blue lines trace sequential NOEs, and pink lines non-sequential inter-residue HN-Hα NOEs supporting α-helical structure.

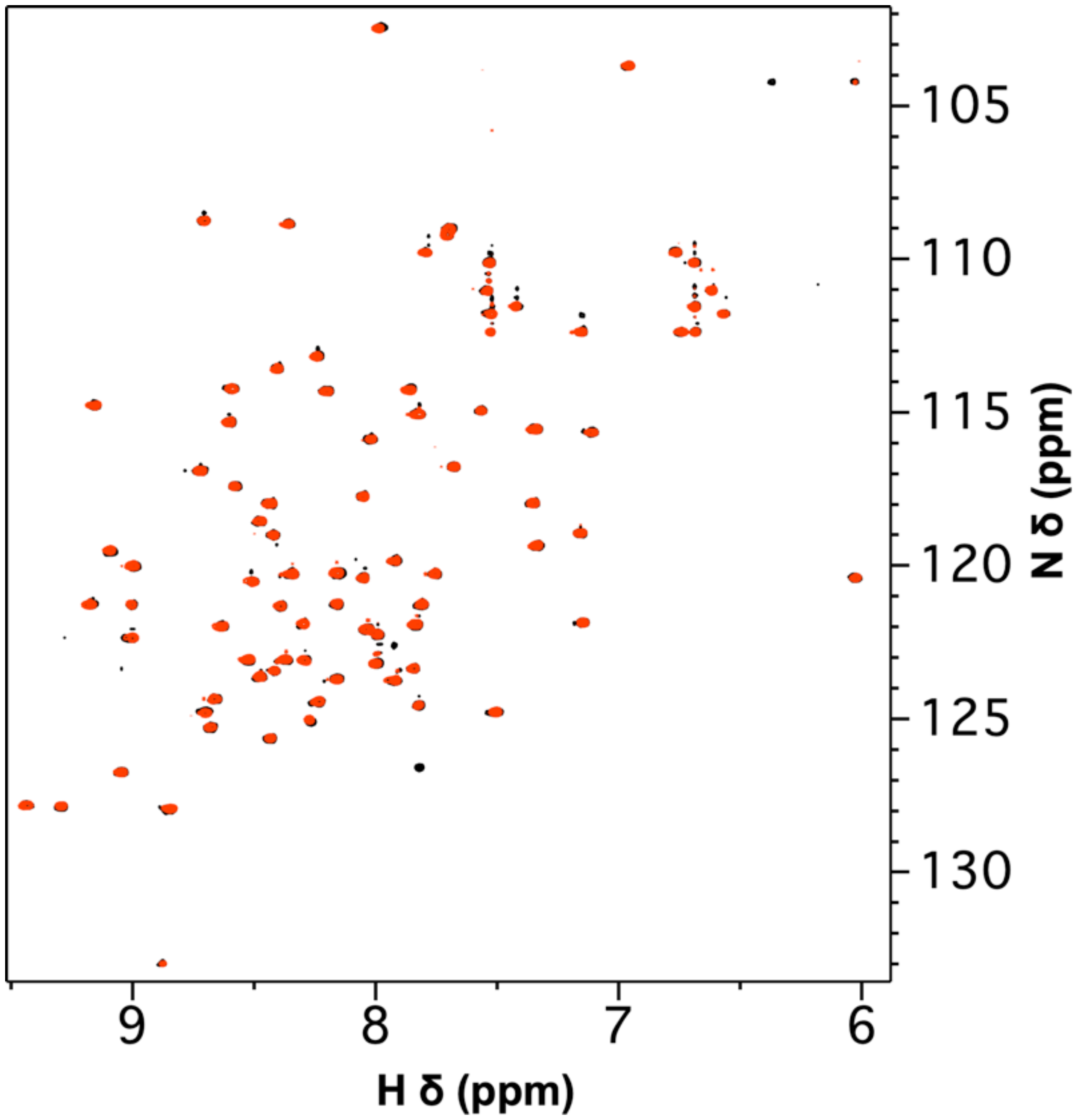


**Supplementary Figure S5 NMR fails to detect interaction between ZC4H2-ZL and ubiquitin.** To a 0.2 mM sample of recombinantly-produced human $^{15}$N-ubiquitin in a buffer containing 150 mM NaCl and 50 mM phosphate, pH 7; we added 0.5, 1.0, 2.0, and 4.0 molar ratios of ZL-$Zn^{2+}$ from a 9.6 mM stock solution. Data were recorded on a Varian Inova 600 MHz spectrometer at a temperature of 25 C. None of the $^{15}$N-ubiquitin $^{1}$H-$^{15}$N HSQC spectra showed changes upon addition of unlabeled ZL-$Zn^{2+}$. The $^{1}$H-$^{15}$N HSQC spectra shown in the figure compare ubiquitin alone (black) to 4:1 ubiquitin: ZL-$Zn^{2+}$ (red). The nearly complete superposition of the two spectra indicates that the two proteins do not interact.

**Table S1 – Statistics for the 20 best NMR structures of the ZL fragment.**

| | | |
|---|---|---|
| NMR restraints (total) | 252 | |
| Distance (total) [a] | 206 | |
| Intraresidue NOE ($\|i-j\| = 0$) | 33 | |
| Sequential NOE ($\|i-j\| = 1$) | 77 | |
| Medium-range NOE ($1<\|i-j\|<5$) | 23 | |
| Long-range NOE ($\|i-j\| \geq 5$) | 49 | |
| Hydrogen bond [b] | 10 | |
| $Zn^{2+}$ coordination restraints [c] | 14 | |
| Dihedral ($\phi$ 19, $\psi$ 19, $\chi_1$ 8) | 46 | |
| *Residual restraint violations* | | |
| Distance (Å) | 0.0471 ± 0.0032 [d] | |
| Dihedral (°) | 1.07 ± 0.14 | |
| *RMS deviations from ideal geometry* | | |
| Bonds (Å) | 0.00467 ± 0.00020 | |
| Angles (°) | 0.75 ± 0.06 | |
| Improper torsions (°) | 2.60 ± 0.31 | |
| *Ramachandran plot PROCHECK statistics for ordered residues* | | |
| Residues in most favored regions | 73.8 % | |
| Residues in allowed regions | 21.2 % | |
| Residues in generously allowed regions | 5.0 % | |
| Residues in disallowed regions | 0.0 % | |
| *Coordinate rms deviations (Å)* | Backbone | All |
| NMR ensemble to mean | 0.69 ± 0.28 | 1.39 ± 0.40 |
| NMR ensemble to mean (ordered: K2-H8, Q10-S25) | 0.60 ± 0.25 | 1.22 ± 0.36 |

[a]The PdbStat program was used to remove redundant and structurally non-informative NOEs from the final restraints described in this table.
[b]Two distance restraints (1.5-2.5 Å for NH-O and 2.5-3.5 Å for N-O) for each of five hydrogen bonds were included to enforce hydrogen bond linearity.
[c]Two restraints per Cys-$Zn^{2+}$ bond (from Sγ and Cβ to $Zn^{2+}$) were used to maintain linearity for a total of eight, and an additional six Sγ-Sγ restraints between each Cys-Cys pair combination were used to enforce tetrahedral geometry.
[d]All values are given as mean ± standard deviation.
